# Energy supply per neuron is constrained by capillary density in the mouse brain

**DOI:** 10.1101/2020.02.03.932434

**Authors:** Lissa Ventura-Antunes, Suzana Herculano-Houzel

## Abstract

Neuronal densities vary enormously across sites within a brain. Does the density of the capillary bed accompany the presumably larger energy requirement of sites with more neurons, or with larger neurons, or is energy supply constrained by a mostly homogeneous capillary bed? Here we find evidence for the latter across various sites in the mouse brain and show that as a result, the ratio of capillary cells per neuron, and thus presumably blood and energy supply per neuron, decreases uniformly with increasing neuronal density and therefore smaller average neuronal size across sites. Additionally, we find that local capillary density is also not correlated with local synapse densities, although there is a small but significant correlation between lower neuronal density (and therefore larger neuronal size) and more synapses per neuron within the restricted range of 6,500-9,500 across cortical sites. Further, local variations in the glial/neuron ratio are also not correlated with local variations in number of synapses per neuron or local synaptic densities. These findings suggest that it is not that larger neurons, neurons with more synapses, or even sites with more synapses *demand* more energy, but simply that larger (and thus fewer) neurons have more energy available per cell, and to its synapses as a whole, than smaller (and thus more numerous) neurons due to competition for limited resources supplied by a capillary bed of fairly homogeneous density throughout the brain.

**Significance Statement:** The brain is an expensive organ and at rest already uses nearly as much energy as during sensory activation. To ultimately determine whether the high energy cost of the brain is driven by an unusually high energy demand by neurons or constrained by capillary density in the organ, we examine whether sites in the mouse brain with more neurons, larger neurons, or more synapses have more capillary supply, and find instead that capillary density is mostly homogeneous across brain structures. We propose that neurons are constrained to using what energy is available, with little evidence for adjustments according to local demand, which explains its high risk of ischemia and vulnerability to states of compromised metabolism, including normal aging.

## Introduction

The human brain ranks second amongst organs in absolute energy cost at rest only to the liver (Rolfe and Brown, 1997; Mink et al., 1981; Aschoff et al., 1971). Such high cost is commonly attributed to synaptic-mediated neuronal activity (Attwell and Laughlin, 2001; Harris et al., 2012; Ames, 2000), and is entirely met by molecules supplied from the blood and provided to neurons through glial cells whose metabolism is coupled to neuronal activity (Magistretti, 2006; Pellerin and Magistretti, 2016). The high energy cost of the brain, liver and heart puts them at high risk of damage by conditions such as aging that compromise metabolism and oxygen supply (Leach et al., 1998). Elucidating what determines the high energetic cost of the brain is central to understanding healthy and abnormal brain function.

Brain bioenergetics researchers typically consider that the high energetic cost of the brain is driven by a steep energetic *requirement* of neurons, due to costs related to synaptic activity and membrane repolarization (Attwell and Laughlin, 2001; Howarth et al., 2010; Harris et al., 2012). However, energetic use can be determined by *demand* only if it is matched dynamically by a non-limiting supply; otherwise, it is constrained by energetic *supply* that limits work. The former is the case of modern cities where electricity supplied to homes is non-limiting, and demand is free to vary, but is always met; the latter is the case in rural homes that already consume most of what little electricity they are supplied, and cannot sustain both an electric shower and air conditioning running simultaneously (Banavar et al., 2002). In biology, this is evident in hummingbirds, forced into torpor when food is insufficient (Hainsworth et al., 1977).

How much the elevated rate of energy use by the brain represents of its energetic supply is a key open question. Importantly, there is reason to suspect that brain energy use at rest is already close to the limits established by supply. First, both local glucose use and blood flow at rest depend linearly on local capillary density in rat (Klein et al., 1986) and monkey (Noda et al., 2002). Task-related variations in local cerebral blood flow (CBF) are very small, in the order of 2%, and of at most 8% in primary sensory areas, such that total CBF remains remarkably constant throughout the day of healthy individuals (Schölvinck et al., 2008; Lin et al., 2010). These small local variations in CBF happen with hardly any capillary recruitment (Kuschinsky and Paulson, 1992; Hudetz, 1997). Additionally, blood flow rates in the resting brain are already as high as first achieved in skeletal muscle during exercise (Madsen et al., 1993; Jie et al., 2014).

Here we examine whether the distribution of capillaries in the brain is consistent with a supply-limiting or a demand-based scenario. To that end, we determine whether capillary density is homogeneous throughout the mouse brain regardless of local variations in local neuronal density, which would be expected if it were determined in a system-wide manner by physical scaling limitations (Banavar et al., 2002); or whether it varies coordinately with local neuronal densities, which would be consistent with demand-based reorganization of the capillary bed in response to use, whether because sites with more neurons have higher energy use simply because of the larger number of neurons, or because larger neurons (in sites of lower local neuronal densities) require more energy individually (Attwell and Laughlin, 2001). To gain insight on the particular issue of energy availability per neuron, we calculate its proxy, the local endothelial-to-neuronal-cell (E/N) ratio, and examine how it varies locally in the mouse brain depending on local neuronal densities. Finally, we examine whether variations in energy availability per neuron might also be related to local variations in numbers of synapses per neuron, by combining our data with direct counts of synapse densities in identified locations in the mouse brain published recently (Zhu et al., 2018).

## Material and Methods

### Ethics Statement

All animal use in this project was approved by the Committee on Ethical Animal Use of the Health Sciences Center (CEUA-CCS), Federal University of Rio de Janeiro (UFRJ) with protocol number 01200.001568/2013-87.

### Experimental design

We used structured illumination confocal imaging to allow quantification of cells and microvasculature in individual sections (2D) and stacks (3D) of multiple locations in the brain of five adult mice. In two of those mice, subjected to detailed 3D quantification, brain microvasculature was revealed by injection of a fluorescent tracer (FITC-Dextran, see below) into the caudal vein; in the other three, subjected to much more time-efficient 2D quantification, brain capillaries were revealed by immunohistochemistry against collagen IV, which labels the basal lamina of blood vessels, once we established that measurements of capillary density were indistinguishable between the two labeling methods (see below). Immunohistochemistry against the neuronal marker NeuN (Mullen et al., 1992) was used to reveal neurons; glial cells were identified by exclusion of neurons and capillary-associated cells from the total number of cell nuclei visualized with DAPI.

We quantified capillary area fraction (which is identical to capillary volume fraction) and cell densities in ten easily definable brain structures (cerebral cortical gray and white matter, cerebellar gray and white matter, thalamus, hypothalamus, striatum, hippocampal and cerebellar granular and molecular layers) imaged with structured confocal illumination in a total of 5 mice, comprising 867 stacks analyzed in 3D (two mice, total of 32,270 cells) and 750 ROI acquired and analyzed in 2D (three mice, total of 196,206 cells; Table 1). While image stacks allow detailed visualization of microvasculature in 3D, two-dimensional analysis of image montages allows for larger sites with larger cell populations to be sampled in less time (Fig. 1). We thus compared the two approaches in this study to determine whether 2D analysis of the distribution of capillaries and neurons are a viable and more efficient alternative to 3D quantification.

**Table 1:** Numbers of cells, stacks and ROIs analyzed in each of ten structures of interest in the mouse brain. CerCx, cerebral cortex; GM, gray matter; WM, white matter; GL, granular cell layer; ML, molecular cell layer.

| Structure | Mouse 01 | Mouse 02 | Mouse 03 | Mouse 04 | Mouse 05 |
| --- | --- | --- | --- | --- | --- |
| **CerCx, GM** | 32 stacks<br>2,516 cells | 54 stacks<br>3,495 cells | 76 ROI<br>63,022 cells | 57 ROI<br>17,613 cells | 34 ROI<br>12,053 cells |
| **CerCx, WM** | 60 stacks<br>1,497 cells | 69 stacks<br>2,531 cells | 59 ROI<br>2,651 cells | 19 ROI<br>1,636 cells | 27 ROI<br>3,931 cells |
| **Thalamus** | 83 stacks<br>4,349 cells | 42 stacks<br>1,997 cells | 21 ROI<br>9,096 cells | 15 ROI<br>5,442 cells | 16 ROI<br>3,634 cells |
| **Hypothalamus** | 45 stacks<br>2,816 cells | 34 stacks<br>1,861 cells | 13 ROI<br>5,358 cells | 12 ROI<br>2,260 cells | 11 ROI<br>2,125 cells |
| **Striatum** | 59 stacks<br>3,283 cells | 43 stacks<br>2,693 cells | 29 ROI<br>15,539 cells | 16 ROI<br>6,284 cells | 18 ROI<br>6,467 cells |
| **Hippocampus, ML** | 40 stacks<br>137 cells | 73 stacks<br>202 cells | 9 ROI<br>136 cells | 9 ROI<br>88 cells | 8 ROI<br>118 cells |
| **Hippocampus, GL** | 25 stacks<br>470 cells | 21 stacks<br>368 cells | 13 ROI<br>726 cells | 13 ROI<br>310 cells | 9 ROI<br>609 cells |
| **Cerebellum, GL** | 37 stacks<br>2,105 cells | 23 stacks<br>1,167 cells | 43 ROI<br>16,850 cells | 22 ROI<br>4,084 cells | 34 ROI<br>5,892 cells |
| **Cerebellum, ML** | 41 stacks<br>361 cells | 26 stacks<br>179 cells | 42 ROI<br>3,982 cells | 23 ROI<br>1,313 cells | 35 ROI<br>1,632 cells |
| **Cerebellum, WM** | 32 stacks<br>138 cells | 28 stacks<br>105 cells | 29 ROI<br>1,746 cells | 15 ROI<br>778 cells | 23 ROI<br>831 cells |
| **Total** | 454 stacks<br>17,672 cells | 413 stacks<br>14,598 cells | 334 ROI<br>119,106 cells | 201 ROI<br>39,808 cells | 215 ROI<br>37,292 cells |

**Figure 1.**
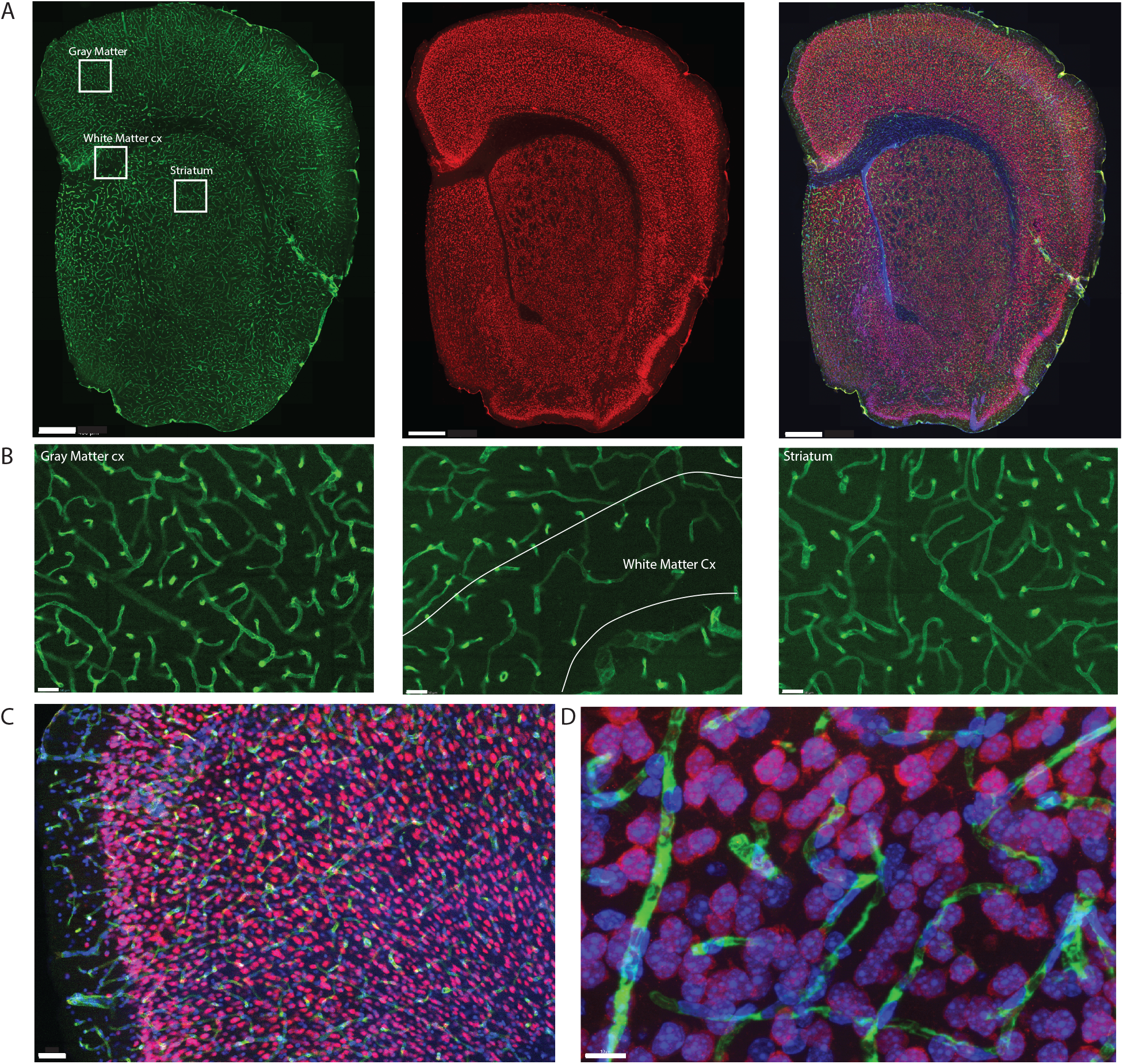
Example of triple labeling for vasculature, neurons, and all cell nuclei. Shown is a coronal section of a mouse brain label by systemic injection of FITC-Dextran. A. Coronal section of mouse brain showing microvasculature labeled with injection of FITC-dextran (left, green), neuronal nuclei labeled with NeuN-Cy3 (center, red), and all nuclei labeled with DAPI (right, blue in merged images). Scale bar, 400μm. **B.** Zoom of insets in A using the 63x objective with the thinner depth of field illustrating the extent of differences in capillary densities across brain sites. Scale bar, 50 μm. **C.** Section through cortical gray matter under 20x magnification stained to reveal Collagen IV (green), NeuN (red) and cell nuclei (blue). Scale bar, 50 μm. **D.** Maximal projection image of 3D stack through cortical gray matter imaged under 63x magnification. Scale bar, 10μm.

From all DAPI-labeled cell nuclei in each stack or 2D ROI, we identified all NeuN-positive cell nuclei as neurons; all nuclei directly associated with collagen IV-labeled or FITC-dextran filled capillaries as endothelial and associated cells (which includes eventual pericytes; heretofore, “endothelial cells”), and by elimination, all remaining nuclei were deemed glial cells (Fig. 2). Cell densities were calculated as cell nuclei per mm^3^ or mm^2^, depending on the stack volume or ROI area analyzed to include capillaries only. All results are shown side by side in 2D and 3D analyses.

**Figure 2.**
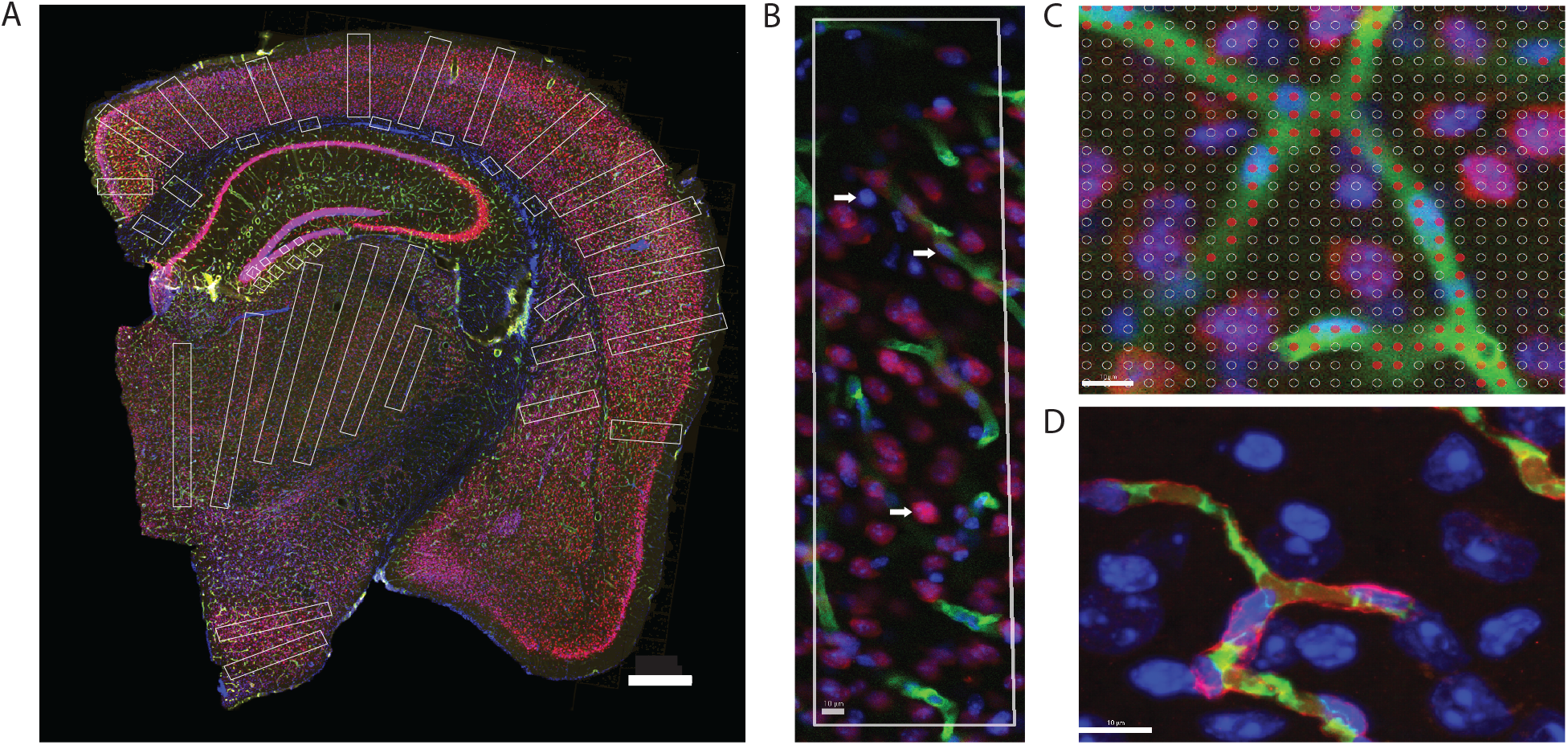
Two-dimensional image analysis. **A.** Areas sampled for 2D quantification are shown in white across structures in a coronal section of the mouse brain. Scale bar, 500 μm. **B.** ROI in the cortical gray matter showing microvasculature labeled with anti-collagen IV (green), neurons labeled with anti-NeuN (red) and Dapi to reveal all cell nuclei (blue). Cell nuclei appearing inside green-labeled blood vessels are considered capillary-associated nuclei; cell nuclei double-labeled in red are considered neuronal cell nuclei; all remaining nuclei are identified by exclusion as glial cell nuclei. **C.** A Cavalieri estimator grid of 4 μm spacing (shown) was applied to images acquired under 20x magnification, and a 2μm grid was applied to images acquired under 63x magnification. The points marked in red correspond to the area of the image occupied by capillaries. Scale bar, 10μm **D**. Superposition of FITC-dextran (green, lumen) and collagen IV (red, basal lamina) labeling of capillaries. Single image acquired under 63x magnification. Scale bar, 10μm.

### Experimental animals

Five male Swiss mice aged 2.5 months were analyzed. Two mice received an injection of 200 mg/kg of fluorescein-isothiocyanate-dextran (FITC-dextran 70 kDa) in the caudal vein, to label the lumen of all blood vessels (Fig. 2). One hour after the injection, the half-life required for distribution in the lumen of blood vessels (Wang et al., 2016), the animals were deeply anesthetized with Xylazine (15 mg/Kg) and Ketamine (100 mg/Kg), decapitated, and their brain was quickly removed and fixed by immersion in 4% of paraformaldehyde in phosphate buffer for one hour followed by 30% sucrose in 0.1 M phosphate buffer for cryoprotection. Using a Leica microtome, the cerebellum was cut into a series of 40 μm thick sagittal sections, and the remaining tissue was cut into a series of 40 μm coronal sections. All sections were stored at −20°C in antifreeze solution (30% polyethylene glycol and 30% glycerol in 0.1 M phosphate buffer) until further use. Three other mice were similarly deeply anesthetized with Xylazine and Ketamine and perfused through the heart with 0.9% saline followed by 4% paraformaldehyde in phosphate buffer. The brain was removed and post-fixed for one hour by immersion in 4% of paraformaldehyde in phosphate buffer, then sectioned as above.

### Immunofluorescence

One in every six sections of each brain was subjected to immunohistochemistry and counterstaining with DAPI. Anti-NeuN antibody was used to reveal neurons; additionally, those sections from the three animals not injected with FITC-Dextran were subjected to immunohistochemistry against collagen IV to reveal cell nuclei intimately associated with capillaries (Fig.2)

Each section was washed in phosphate buffer (PB) for five minutes, heat for one hour at 70 °C in a 0.1 M solution of boric acid, pH 9.0, then incubated for one hour in PB containing 3% normal goat serum (Sigma-Aldrich, LG9023) and 2% bovine serum albumin (Sigma-Aldrich-A2058). Sections from animals not injected with FITC-Dextran were next incubated under stirring for 48 hours at 4°C in PB containing rabbit polyclonal anti-collagen IV antibody (Abcam, Ab6586) at a 1:500 dilution. Each section was then washed three times in PB for 5 minutes each and incubated for 2 hours at 4°C in a 1:500 dilution of Alexa Fluor 488 goat anti-rabbit secondary antibody (Abcam, Ab150077). From this step on, all brain sections were treated similarly: washed three times in PB and incubated for 24 hours at 4°C under continuous stirring with rabbit primary polyclonal antibody against Cy3-conjugated NeuN (Millipore, ABN78C3) diluted 1:1000 in PB. All sections were then labeled with DAPI (4’,6-Diamidino-2-Phenylindole Dilactate, from stock solution at 20 mg/l) to provide counter-staining for identification of brain structures and allow visualization of all cell nuclei in each section. The sections were then mounted on glass slides and coated with Vectashield (Vector Labs, Cat. number H-1400) for viewing under fluorescence microscopy.

### Structures analyzed

The ten structures of interest targeted for analysis (gray matter (GM) and white matter (WM) of the cerebral cortex; striatum; thalamus; hypothalamus; granular and molecular layers of the hippocampal dentate gyrus; white matter and granular and molecular layers of the cerebellum) were defined according to the Mouse Brain Atlas (Franklin and Paxinos, 1997).

### Microscopy

Two Zeiss AxioImager M2 microscopes equipped with an Apotome 2 for confocal imaging under structured illumination (Carl Zeiss, Jena, Germany) and driven by StereoInvestigator software (Microbrightfield Bioscience, Williston, VT) were used for all image acquisition.

We first delineated the boundaries of the ten brain structures of interest in each section. For each structure, image stacks for 3D analysis were acquired using systematic random sampling in Stereo Investigator under magnification with a 63x objective (Plan-ApoChromat), which gives a depth of field of 0.5 μm with oil, with an optical plane of 0.5 μm and step size of 0.5 μm. Stacks were 1388 ×1040 μm wide × 30 μm thick and typically contained 60 images each, spanning most of the full mounted thickness of the section. Within each stack, 3D counting probes placed were variable in size according to the structure (Table 2) and were placed to exclude the sectioned surfaces and zones outside the structure of interest. The grey matter of the cerebellar cortex was imaged as a whole and analyzed separately into molecular and granular layers. For 2D analysis, image composites of the entire brain sections were acquired under 20x magnification (EC Plan-NeoFluar/420350-9900), which gives a thicker depth of field of ca. 4 μm, and counting probes consisted of rectangular regions of interest (ROIs) placed to cover large areas of the target structures in each brain section (see Fig. 2). In the cerebral cortex, rectangular probes spanned the entire grey matter from pia to the white matter border, and 5-15 probes of ca. 800 μm width were placed at regular intervals along each section. For all other structures, 2-8 rectangular probes were placed in each section, seeking to maximize the sampled surface of each structure (Fig. 2A).

**Table 2:** Size of 3D counting probes per structure. CerCx, cerebral cortex; GM, gray matter; WM, white matter; GL, granular cell layer; ML, molecular cell layer.

|  |  |
| --- | --- |
| CerCx GM | X 127 y 96 z 20 $\mu\text{m}$ |
| CerCx WM | x 80 y 75 z 20 $\mu\text{m}$ |
| Thalamus | x 100 y 95 z 20 $\mu\text{m}$ |
| Hypothalamus | x 100 y 95 z 20 $\mu\text{m}$ |
| Striatum | x 100 y 95 z 20 $\mu\text{m}$ |
| Hippocampus ML | x 40 y 40 z 20 $\mu\text{m}$ |
| Hippocampus GL | x 30 y 30 z 20 $\mu\text{m}$ |
| Cerebellum GK | x 40 y 40 z 20 $\mu\text{m}$ |
| Cerebellum ML | x 40 y 40 z 20 $\mu\text{m}$ |
| Cerebellum WM | x 40 y 40 z 20 $\mu\text{m}$ |

Although highly time-consuming, detailed analysis of 3D images was performed first to rule out bias in data acquisition due to any preferential organization of capillaries relative to the plane of section, given the three-dimensional nature of the distribution of the capillary network. To ensure feasibility of future analyses of a large number of species, and to allow for a much larger sample size, we compared the results of the 3D analysis of small image stacks with the much faster and more practical analysis of 2D image composites, which permitted the analysis of much larger ROIs and numbers of cells in half as much time. Those analyses had to be performed in separate animals due to bleaching of the immunofluorescence during acquisition of image stacks. Due to software limitations that precluded acquiring large composites at 63x, 2D image composites had to be acquired at 20x, which leads to inflated estimates of area (or volume) fraction occupied by capillaries due to the projection of oblique vessels given the larger thickness of the optical sections at 20x (ca. 4 μm) compared to 63x (ca. 0.5 μm). For the same reason, estimates of local cell densities will also differ between the two methods. However, we hypothesize that all relationships across variables should remain unaffected by the method of image acquisition. All results are thus presented side by side for 3D and 2D analyses. Data points overlap across animals (shown in different symbols in all figures), and are thus analyzed jointly.

### 3D analysis

Stacks in the targeted brain structures of two FITC-injected mice were acquired using systematic random sampling (SRS) of each structure in each of 1 of 6 sections. From each brain, 13 and 18 sagittal sections through the cerebellum and 24 and 23 coronal sections through the remaining brain structures, including the cerebrum, were analyzed. A total of 867 stacks were analyzed across the ten brain structures in two mice (Table 1, mouse 01 and mouse 02).

### 2D analysis

From each of three animals, 21, 12 and 17 sections through the cerebellum and 21, 20 and 14 sections through the remaining brain structures were analyzed. A total of 750 ROIs were imaged and analyzed across the ten brain structures (Table 1, mice 03-05).

### Image analysis

In each stack and ROI analyzed, we manually identified each DAPI-stained cell nucleus in the structure site as neuronal (when it expressed NeuN immunoreactivity), endothelial or belonging to other capillary-associated cells (when the cell was intimately associated with FITC-labeled capillary lumen or collagen IV-stained basal lamina, which includes pericytes), or glial (by exclusion; Fig. 2B).

Microvascular area fraction (hereafter termed “capillary area fraction” for simplicity) was estimated in image stacks and 2D ROIs using the Cavalieri estimator in StereoInvestigator using a grid of points separated by 2 μm in 3D images and 4 μm in 2D images (Figure 2C). Capillary *area* and *volume* fraction are interchangeable (25) and refer to the fraction of tissue formed by endothelial cells and the lumen of the capillaries that they form. In order to restrict analysis to capillaries, where stacks included large vessels, the counting probe was reduced to avoid it, or else the stack was discarded. In 2D, ROIs were placed to avoid large vessels.

Labeling capillaries with FITC-dextran or immunohistochemistry to collagen IV is expected to lead to slightly different measurements of capillary density expressed as area (or volume) fraction, because while the former labels the lumen of blood vessels, the latter labels the basal lamina of the cells that form the walls of the vessels. However, estimates of numbers of capillary-associated cells per site should remain unaffected. To evaluate how much the difference in measured capillary volume fraction depending on staining method would impact our results, one additional 1-in-6 series of brain sections of mouse #01, which received the injection of FITC-Dextran, were double-labeled for collagen IV using the same primary antibody as before, but a different secondary antibody, conjugated to Alexa Fluor 546 (1:500/#A11010 – Life). Fig. 2D illustrates the coincident but non-overlapping double labeling of lumen and basal lamina of the microvasculature and the associated DAPI-stained cell nuclei. Quantification of the area fraction covered by capillaries in 2D composites acquired at 20x showed a detectable but not statistically significant difference between the vascular fraction estimated with collagen IV (7.4±0.3%) compared to FITC-Dextran (6.6±0.4%).

### Correlations with synaptic density

In order to determine whether local densities of synapses correlate with local densities of neurons and capillaries and to establish how variable are the ratios of synapses per neuron and number of synapses supplied per capillary cell, we proceeded with the quantification of four sections from the brain of mouse #4 that matched the respective levels of the four coronal sections through the cerebral cortex of one individual mouse in the synaptome study of Zhu et al. (2018). We used the Mouse Brain Atlas (Franklin and Paxinos, 1997) to place large ROIs in the same structures. 2D composite images acquired under 20x magnification were analyzed to determine the density of cells and vascular fraction of neocortex in our images for comparison with the total number of synapses labeled with either PSD95 or SAP (Zhu et al. 2018).

### Statistical analysis

All analyses were performed in JMP14 PRO, using non-parametric Spearman correlation coefficients to test the correlation between variables, and regression to mathematical functions to determine the type of relationship between variables.

## Results

To guide the interpretation of our findings, the three main possible scenarios are depicted in Figure 3. The top row illustrates the results expected if steady-state capillary density in the adult brain reflected local variations in neuronal density, and the average neuron used similar amounts of energy regardless of its size or location. In this case, local capillary density should be directly proportional to local neuronal density, and energy availability per neuron, indicated by the ratio of endothelial-associated cells per neuron, should remain constant across brain sites. On the other hand, if larger neurons use more energy, then sites with larger neurons and thus lower neuronal densities should have higher E/N (Figure 3, rows 2 and 3). However, this finding could occur in either one of two other scenarios, depending on whether higher E/N where it happens is due to reorganization of the capillary bed reflecting neuronal *demand*, in which case larger capillary densities would be expected in those sites with larger neurons (and therefore lower neuronal densities; row 3), or is imposed by a lack of variation in capillary density across brain sites (row 2). The three particular cases in which numbers of synapses are constant per volume (left), constant per neuron (center) or larger in larger neurons (right) are depicted separately for each scenario and illustrate the ensuing expected variation in energy availability per synapse as well as how energy availability per neuron would vary together with energy availability per synapse or not.

**Figure 3.**
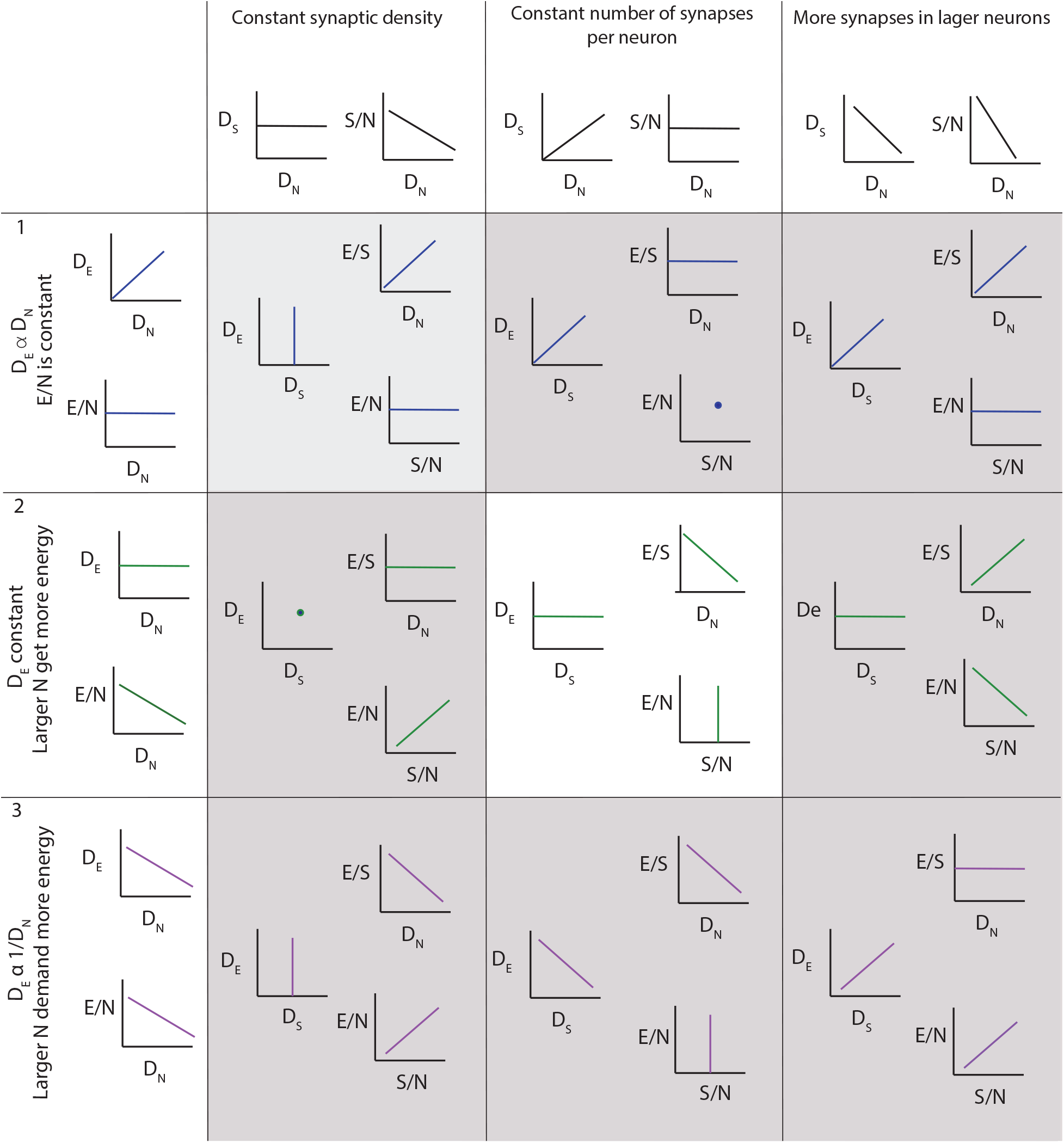
Three of several possible scenarios of relationships among local density of capillary cells (D_E_), local density of neurons (D_N_), energy available per neuron (E/N, estimated as D_E_/D_N_), local density of synapses (D_S_), average number of synapses per neuron (S/N) and average energy supply per synapse (E/S). **Scenario 1 (top row):** neurons demand a certain fixed supply of energy, and the capillary bed adjusts to supply it at a steady-state, such that energy supply per neuron is constant across sites. In this scenario, D_E_ is expected to be proportional to D_N_, and energy available per neuron (E/N) is constant. **Scenario 2 (second row)**: capillary supply to brain tissue is constant across sites, and neurons compete for whatever energy is available. In this case, D_E_ is expected to be relatively constant regardless of local D_N_; as a result, E/N is larger in sites of lower neuronal densities (indicative of larger neurons), where fewer neurons compete for a limited supply of energy. This is the scenario supported by our findings, but incompatible with current models. **Scenario 3 (third row)**: larger neurons demand more energy, and the capillary bed adjusts to supply it, such that energy supply per neuron is larger where neurons are also larger, and thus occur at lower D_N_. This scenario is compatible with current models, but is not supported by our present findings. **Columns** indicate, for each scenario of capillary and neuronal distribution, the expected findings regarding variations across the variables measurable in this study depending on whether **(left column)** local synaptic densities are mostly constant across sites, such that there are more synapses per neuron where neurons are larger (and D_N_ lower); **(center column)** the number of synapses per neuron is mostly constant, such that there are more synapses where there also are more neurons; and **(right column)** larger neurons have many more synapses than smaller neurons, such that local synaptic densities are larger where neurons are larger (and D_N_ lower). Our findings support a constant density of capillaries with fairly constant densities of synapses per neuron, such that energy supply both per neuron and per synapse, and thus energy availability per neuron and per synapse, decreases the higher the local neuronal density, that is, the smaller the average local volume of individual neuronal cells. This is the scenario highlighted in the center panel.

Across the ten brain structures examined, we find that the capillary area fraction is very small, amounting to no more than 3% of the tissue in any structure in 3D image stacks. While individual measurements of local capillary fraction vary 10x-fold across all brain sites, the average capillary fraction per structure varies only about 2-fold across structures, with 95% of measurements ranging around 1.29 ± 1.39% (Table 3). The average capillary fraction is highest in the gray matter of the cerebral cortex (2.22±0.11%), and lowest in the granular layer of the dentate gyrus and subcortical white matter (0.70±0.07%; Table 3). In 2D images, we found similarly small variation in capillary fraction, as well as lower capillary fractions in white matter than in gray matter, despite the artificially higher nominal capillary fractions measured with this method due to the larger optical thickness of the images (Table 4). The capillary fractions we report in 3D analyses are similar to vascular fractions between 2% and 6% reported in the mouse cerebral cortical grey matter (Tsai et al., 2009) and between 2.1% and 3.1% in the cat, macaque and human gray matter cortex (Boero et al., 1999; Pawlik et al., 1981; Lauwers et al., 2008).

**Table 3.** Fractional cellular composition of different structures of the mouse brain, quantified in 3D. Values correspond to the average across sites in each structure of two mice (#1 and #2). CerCx, cerebral cortex; GM, gray matter; WM, white matter; GL, granular cell layer; ML, molecular cell layer; Cb, cerebellum.

| 3D | % capillary fraction | % endothelial cells | % neurons | % glial cells | Glial cells % of non-neuronal cells |
| --- | --- | --- | --- | --- | --- |
| CerCx, GM | 2.22 ± 0.11% | 13.90 ± 0.79% | 57.86 ± 1.53% | 28.22 ± 1.23% | 66.64 ± 1.49% |
| CerCx, WM | 0.75 ± 0.03% | 7.62 ± 0.60% | 5.75 ± 1.15% | 82.61 ± 1.41% | 91.37 ± 0.76% |
| Striatum | 1.43 ± 0.06% | 7.94 ± 0.34% | 62.47 ± 1.10% | 29.57 ± 1.10% | 77.58 ± 0.99% |
| Thalamus | 1.35 ± 0.05% | 10.74 ± 0.52% | 51.24 ± 1.31% | 38.01 ± 1.2% | 77.31 ± 1.00% |
| Hypothalamus | 1.34 ± 0.06% | 10.14 ± 0.69% | 60.62 ± 1.61% | 29.23 ± 1.36% | 73.34 ± 1.64% |
| DG, GL | 0.70 ± 0.07% | 1.46 ± 0.47% | 94.96 ± 1.08% | 3.57 ± 0.92% | 67.46 ± 9.30% |
| DG, ML | 1.10 ± 0.06% | 29.18 ± 3.21% | 10.33 ± 3.29% | 60.48 ± 3.29% | 68.23 ± 3.35% |
| Cb, GL | 1.56 ± 0.06% | 2.58 ± 0.33% | 95.44 ± 0.46% | 1.96 ± 0.33% | 38.86 ± 5.06% |
| Cb, ML | 1.69 ± 0.09% | 13.92 ± 3.76% | 59.77 ± 3.90% | 26.30 ± 3.76% | 54.35 ± 4.93% |
| Cb, WM | 1.25 ± 0.06% | 14.14 ± 2.68% | 0.73 ± 0.52% | 85.12 ± 2.76% | 85.64 ± 2.69% |

**Table 4.** Fractional cellular composition of different structures of the mouse brain, quantified in 2D. Values correspond to the average across sites in each brain structure of three mice (#3, #4 and #5). *Note that the capillary fraction appears higher here than when measured in 3D due to the deeper focal thickness under the lower magnification used for 2D image acquisition, which artificially inflates the relative capillary fraction. Other values are similar between 2D and 3D estimates.

| 2D | % capillary fraction* | % endothelial cells | % neurons | % glial cells | Glial cells % of non-neuronal cells |
| --- | --- | --- | --- | --- | --- |
| CerCx, GM | 8.65 ± 0.21% | 14.37 ± 0.23% | 60.86 ± 0.39% | 24.75 ± 0.36% | 63.06 ± 0.53% |
| CerCx, WM | 5.35 ± 0.27% | 10.63 ± 0.55% | 2.99 ± 0.93% | 86.37 ± 1.18% | 88.55 ± 0.75% |
| Striatum | 7.74 ± 0.25% | 11.88 ± 0.37% | 63.59 ± 0.83% | 24.51 ± 0.71% | 67.14 ± 0.88% |
| Thalamus | 7.58 ± 0.71% | 14.53 ± 0.71% | 45.88 ± 1.89% | 39.57 ± 1.90% | 72.19 ± 1.30% |
| Hypothalamus | 7.28 ± 0.39% | 12.38 ± 0.63% | 57.13 ± 1.60% | 30.47 ± 1.34% | 70.77 ± 1.40% |
| **DG, GL** | 6.17 ± 0.06% | 4.55 ± 0.72% | 90.89 ± 1.46% | 4.55 ± 0.89% | 46.32 ± 5.72% |
| **DG, ML** | 10.91 ± 0.91% | 30.67 ± 1.70% | 13.17 ± 2.34% | 56.15 ± 2.34% | 64.43 ± 1.93% |
| **Cb, GL** | 6.76 ± 0.26% | 3.02 ± 0.17% | 91.00 ± 0.47% | 5.97 ± 0.38% | 63.22 ± 1.83% |
| **Cb, ML** | 7.20 ± 0.25% | 18.77 ± 0.62% | 47.02 ± 1.57% | 34.20 ± 1.48% | 62.47 ± 1.48% |
| **Cb, WM** | 4.16 ± 0.22% | 13.50 ± 0.77% | 2.34 ± 0.56% | 84.14 ± 0.95% | 86.13 ± 0.79% |

Interestingly, capillary-associated (“endothelial”) cells typically constitute between 7 and 14% of all cells forming the brain structures examined (Tables 3, 4), a much larger percentage than could be expected from the average vascular fraction of 0.7-2.2% of the 3D volume that they occupy. This discrepancy indicates that endothelial cells are, on average, much smaller than neurons and glial cells in the tissue. Still, glial cells are consistently the majority (at least 60-70%) of non-neuronal cells in all structures (Tables 3, 4). As expected from our previous studies (reviewed in Herculano-Houzel et al., 2014), the percentage of cells that are neurons is highly variable across structures and sites within a structure; neurons are the vast majority of all cells in the granular layers of the dentate gyrus and cerebellum, a smaller percentage of all cells in other gray matter structures, and rare (but present) within the subcortical white matter of both cerebral and cerebellar cortices (Tables 3, 4).

As expected for two different measurements of the same feature (capillary density), local endothelial cell density correlates well with local capillary area or volume fraction within and across the different brain structures, as well as across animals, especially in the 2D sample, which included a much larger number of sites and cells (Fig. 4A; Table 1). Both capillary fraction and local endothelial cell density vary by a single order of magnitude across sites in the mouse brain, whether measured in 2D or 3D (Fig. 4A). The two capillary density-related variables are correlated within each structure individually (Table 5), and there is good overlap in data points across structures and animals, especially in the larger 2D dataset (Fig. 4A, right), where the overlapping power relationships (Table 5) indicate that the relationship between capillary fraction and cellular composition of capillaries is shared throughout the brain. This agreement suggests that endothelial cell density is a good proxy for the resting rates of blood flow and glucose use (Klein et al., 1986; Noda et al., 2002).

**Table 5.**
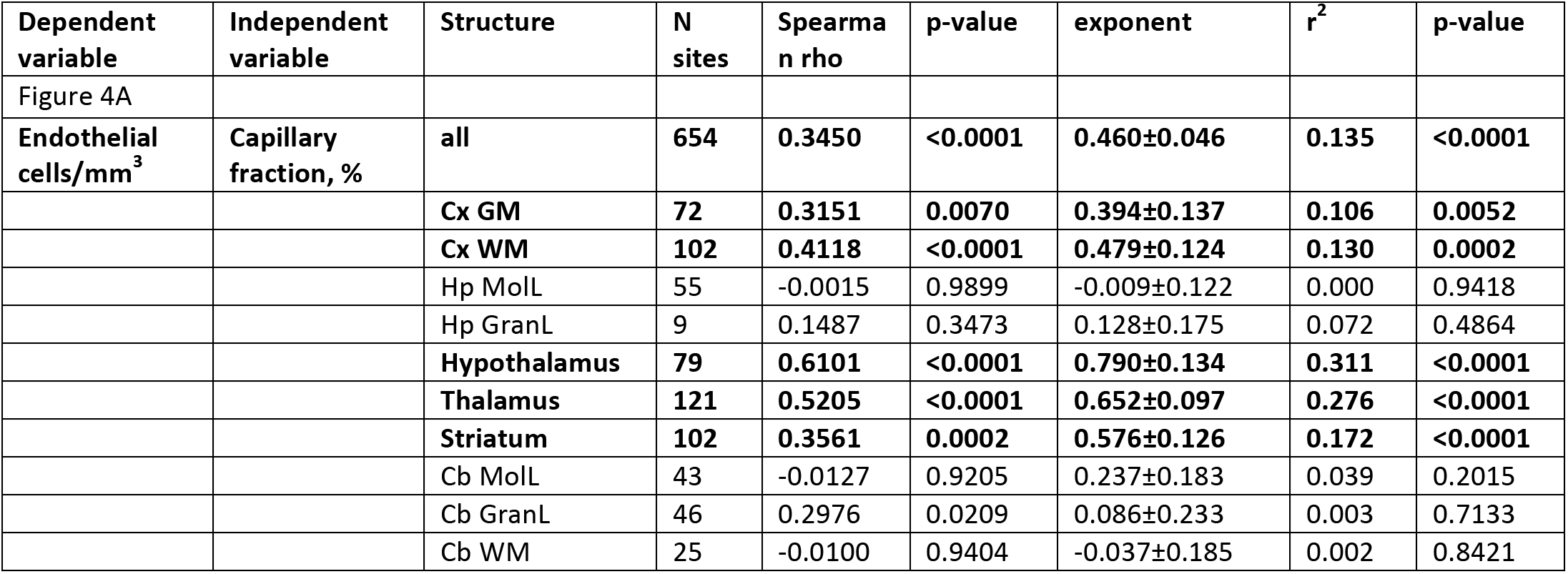
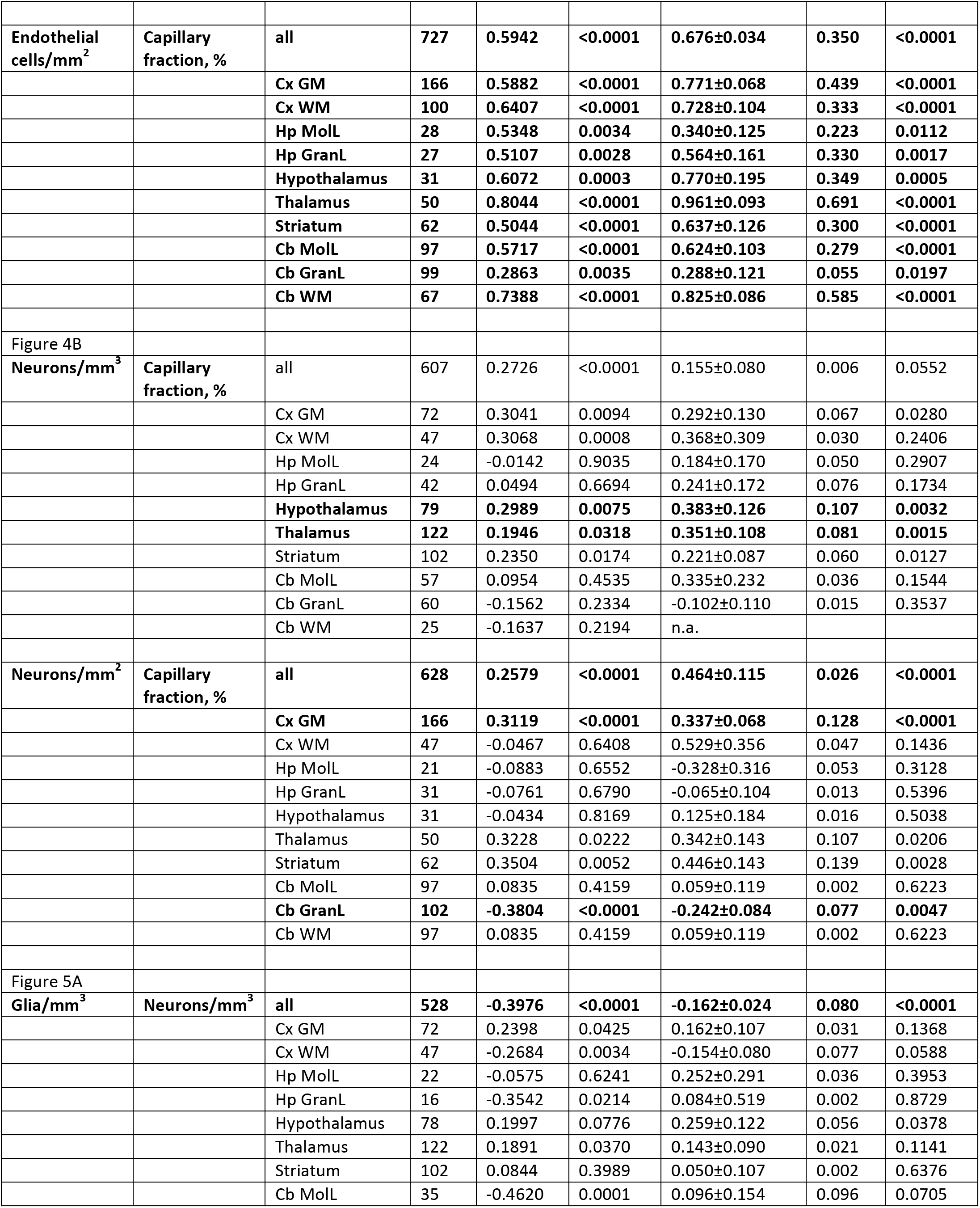
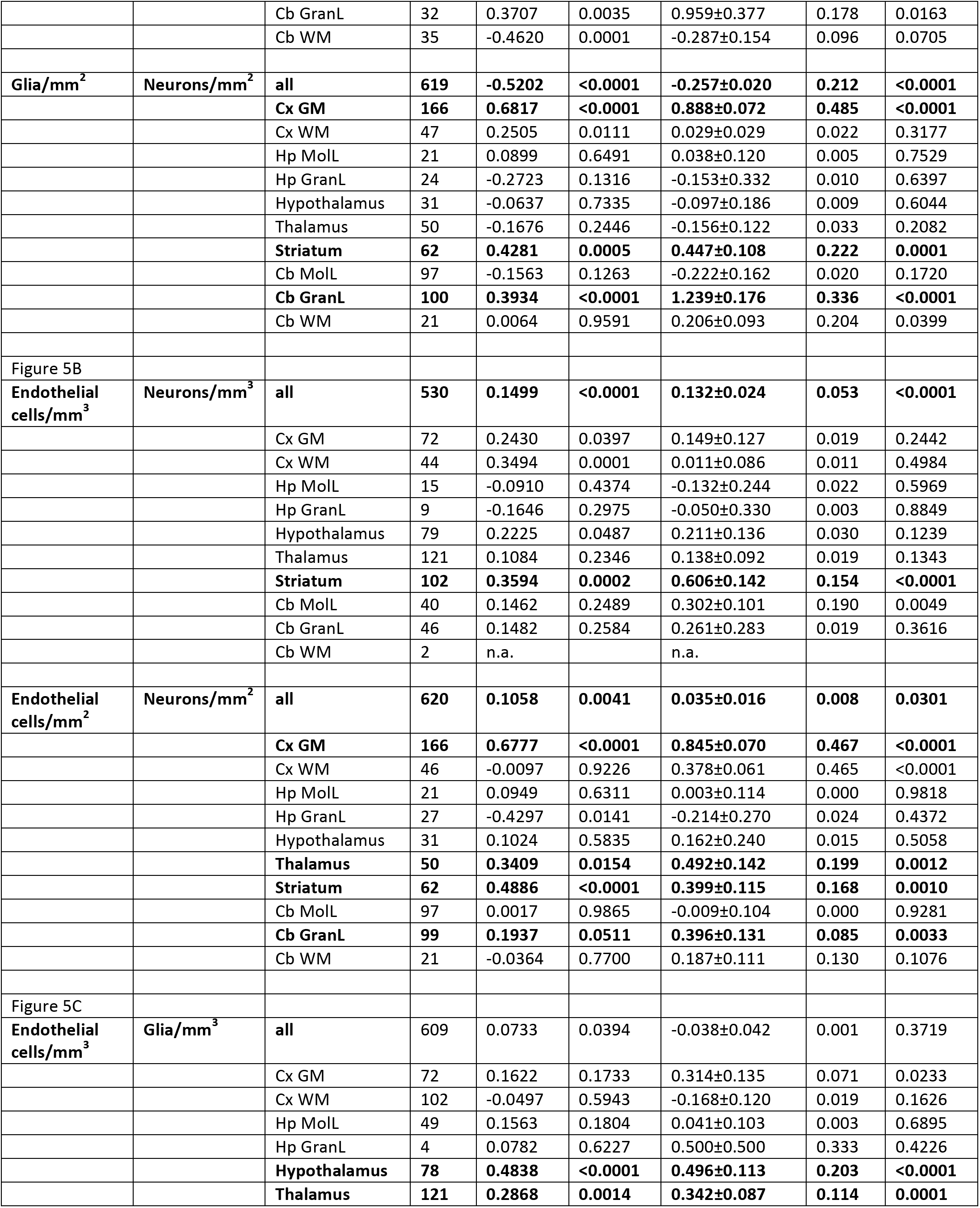
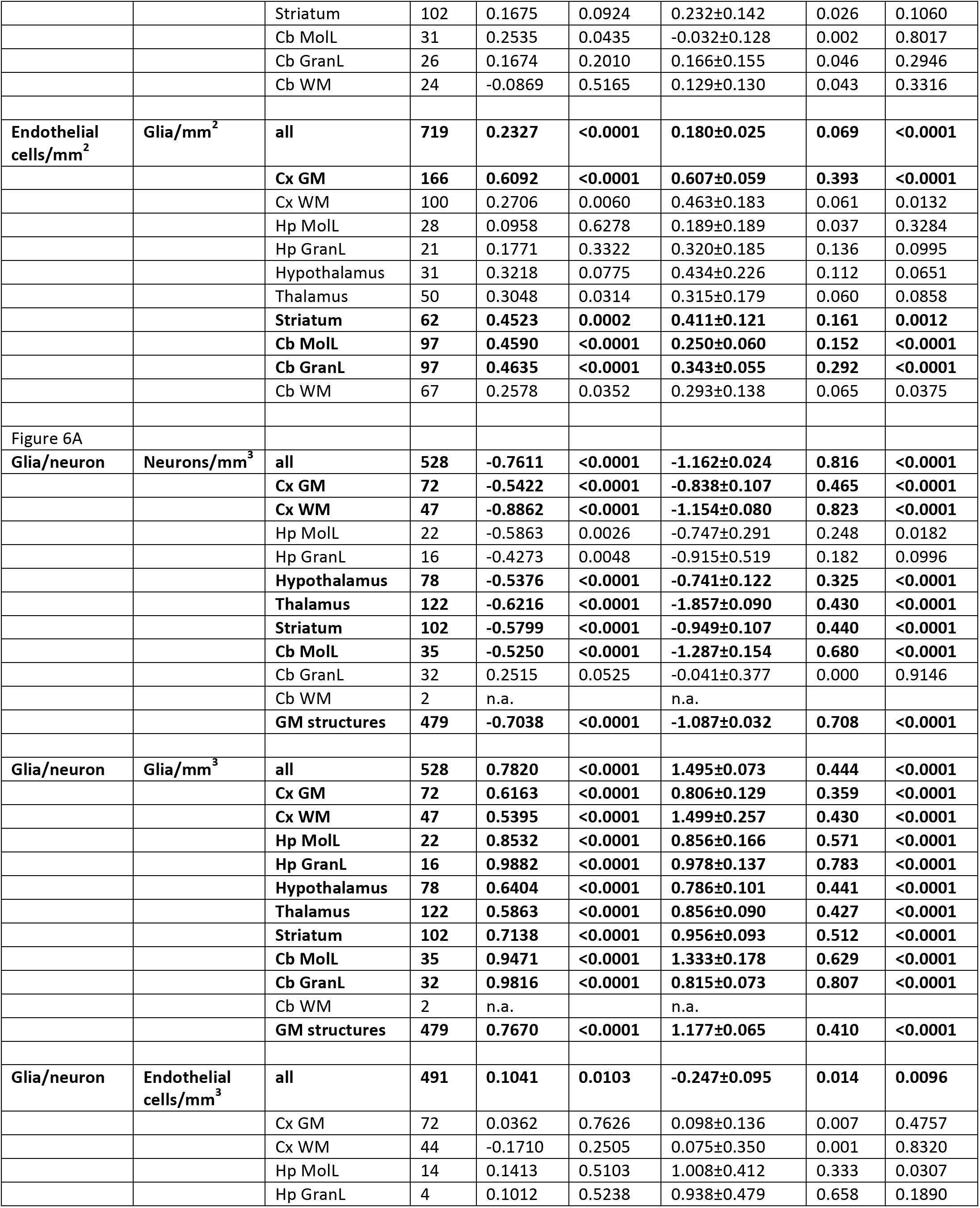
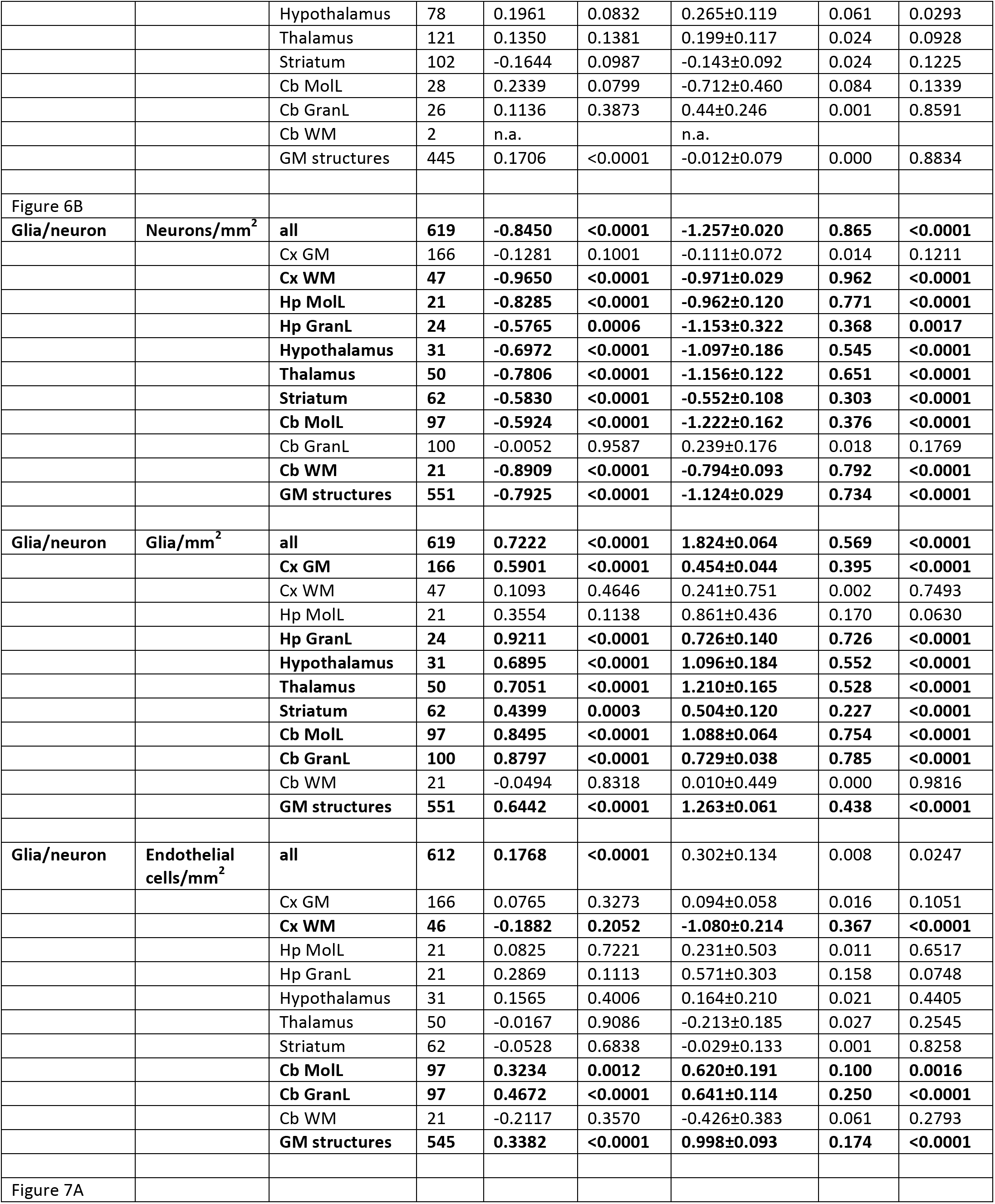
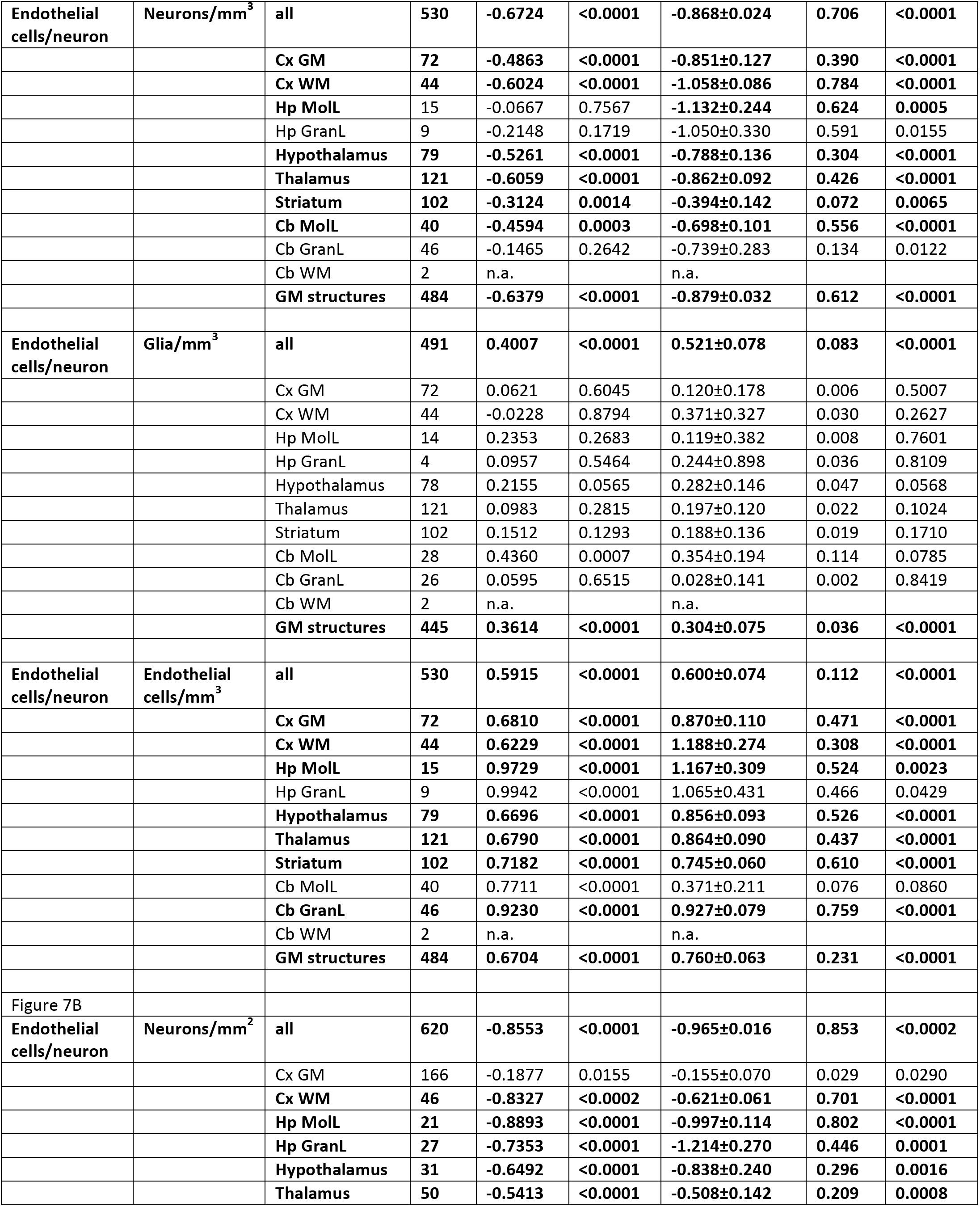
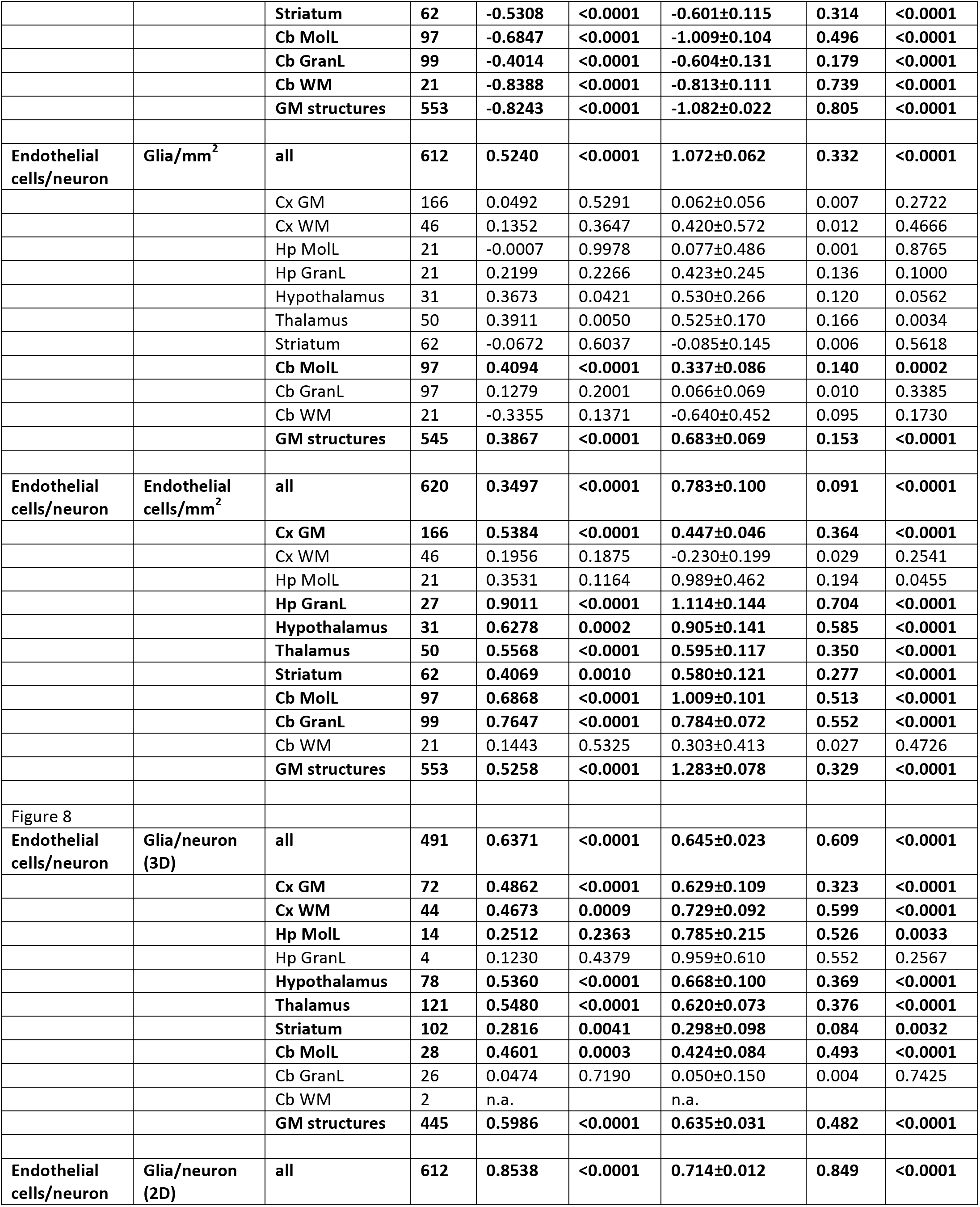
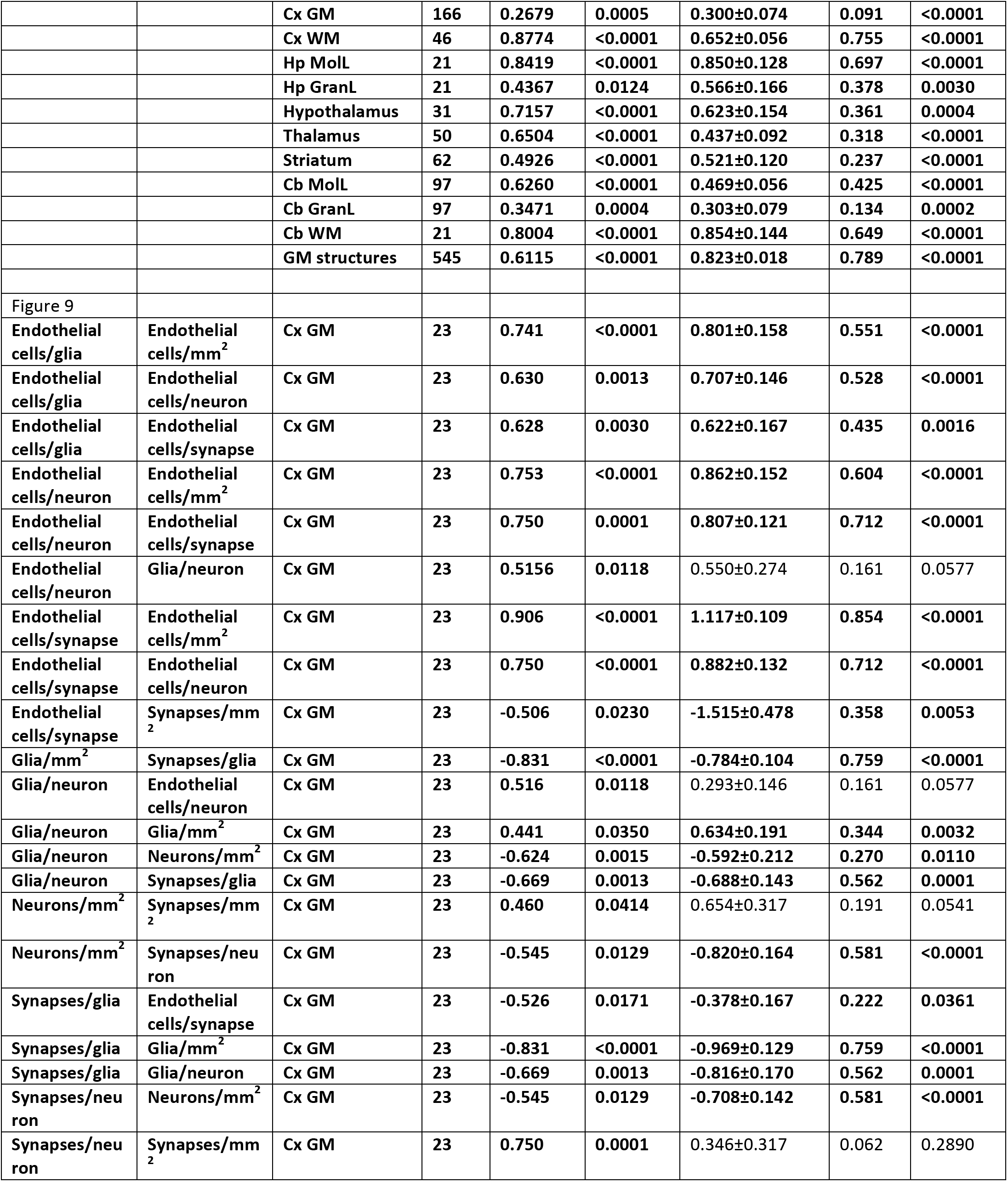
Correlation analysis between variables across sites and animals.

**Figure 4:**
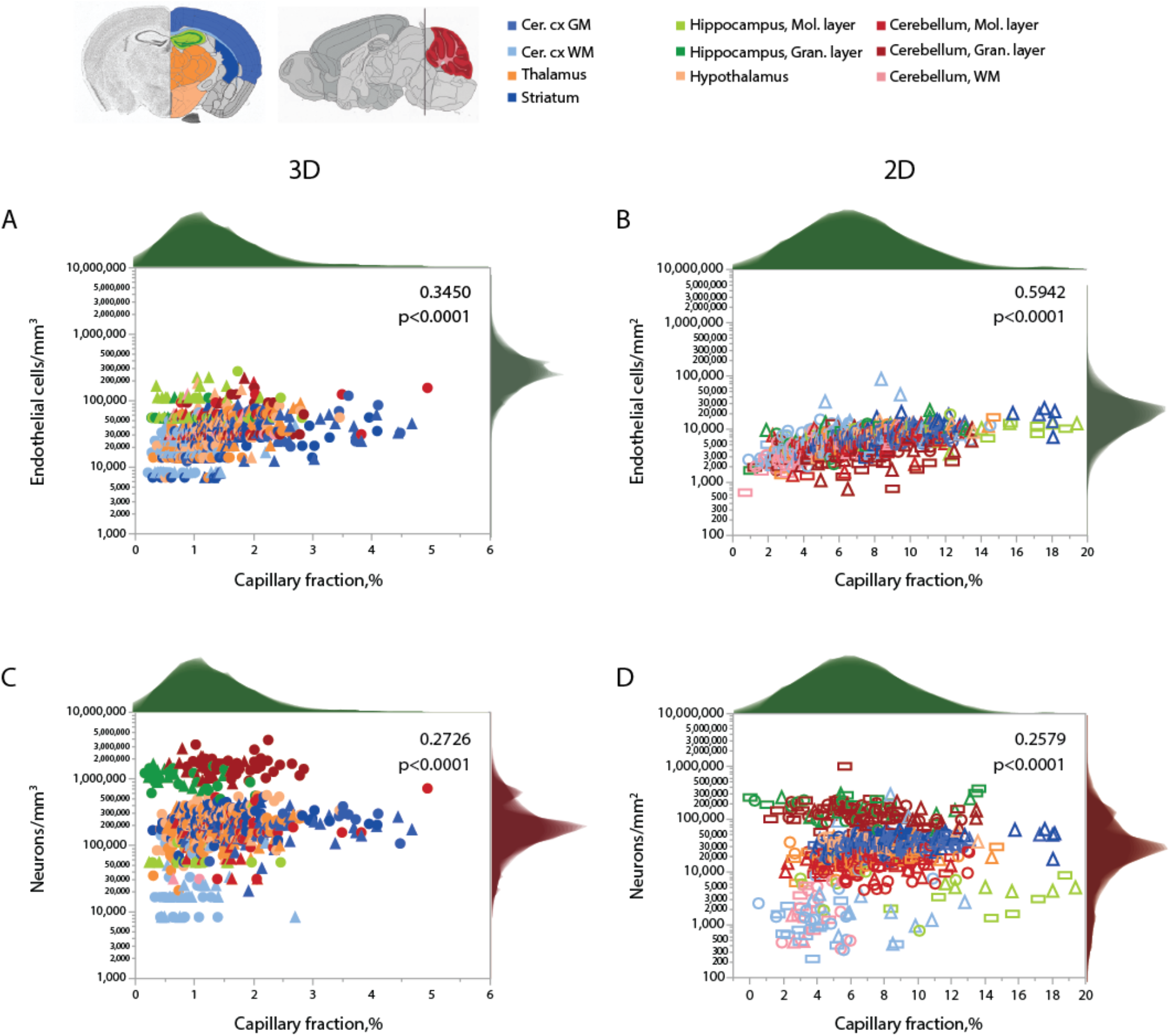
Local capillary fraction correlates with local endothelial cell density, but not with neuronal cell density. **A, B:** Local density of endothelial cells correlates with local vascular fraction across all sites and structures analyzed in 3D (**A**; two animals, shown in circles or triangles, left) and in 2D (**B**; three animals, shown in circles, triangles, or rectangles, right) with the Spearman correlation coefficients and p-values across all animals and sites indicated in each graph. **C, D:** In contrast, local neuronal density does not correlate significantly with local vascular fraction across brain sites and structures in 3D (**C**) or in 2D (**D**). Histograms above and to the right reflect the distribution of all data points in each axis. Graphs in **A/C** and **B/D** are shown with similar X and Y scales for comparison. Correlation coefficients and power exponents for each structure are given in Table 5.

Neuronal densities, in contrast, vary over two orders of magnitude across sites in gray matter structures, from 31,250 to 3,843,750/mm^3^ (Fig. 4B), and are not significantly correlated with local capillary fraction (Figure 4B) across brain sites and structures. In fact, brain structures with enormously different neuronal densities have capillary fractions in the same restricted range (Fig. 4B), and correlations, where significant, can be positive (cortical GM) or negative (cerebellar granular layer; see Table 5 at end of document).

Like endothelial cell densities, variation in glial cell densities is restricted to within one order of magnitude across sites in the mouse brain (Fig. 5A). Within the cortical gray matter, local glial cell densities vary significantly in positive correlation with local neuronal density, as previously observed across sites within both human (Ribeiro et al., 2013) and mouse (Herculano-Houzel et al., 2013) cortices (Fig. 5A; Table 5). In contrast, there is no universal correlation between local neuronal and glial cell densities across brain structures; if anything, there is a modest negative correlation across all sites (Fig. 5A), which is due to the slightly more elevated glial cell densities in white matter sites.

**Figure 5.**
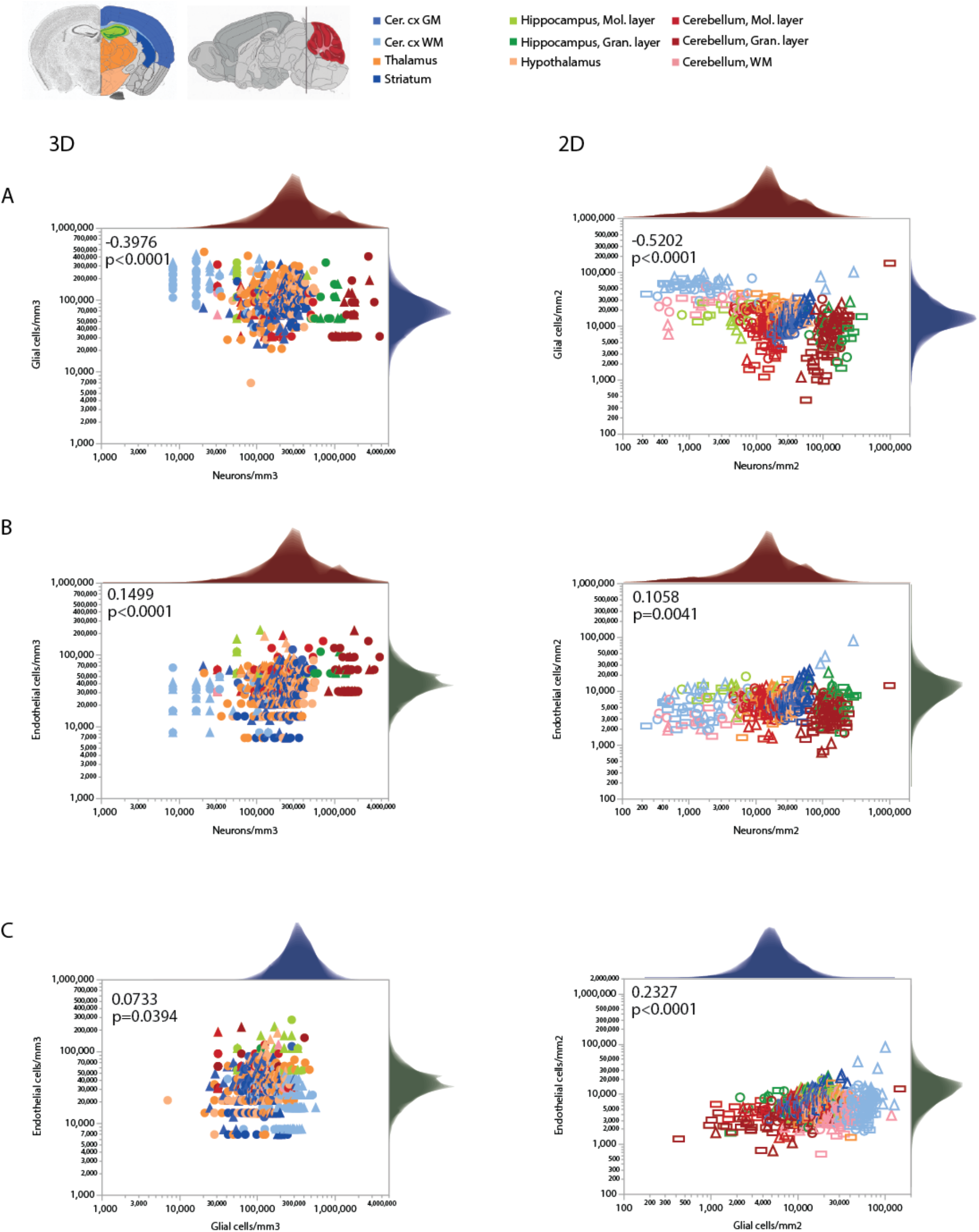
Local densities of different cell types vary concertedly only within some brain structures. Each plot is accompanied by the respective distribution histograms aligned with the X and Y axes. Spearman correlation coefficients and p-values across all structures and sites are indicated in each graph. **A.** Glial cell densities span just over one order of magnitude across structures, while neuronal cell densities span two orders of magnitude. While the correlation across all data points is significant and negative in both 3D and 2D datasets, analysis at that level ignores obvious differences across structures (see Table 4). **B.** Similarly, endothelial cell densities are concentrated within one order of magnitude across structures and sites, and are similar across structures with very different neuronal densities. Still, local endothelial and neuronal cell densities are significantly and positively correlated within the cortical gray matter, the thalamus, and the striatum (see Table 4). **C.** Glial and endothelial cell densities, which vary little across sites and structures in comparison to neuronal densities, are not strongly correlated across brain structures, and weakly correlated within some (see Table 5). All graphs are shown with similar X and Y scales for comparison.

Importantly, a large range of neuronal densities occurs with fairly similar endothelial cell densities across structures in the mouse brain, with only a weak overall trend toward higher endothelial cell densities where neuronal densities are higher (Fig. 5B). A significant increase in local endothelial cell density in sites with increasing local neuronal density is found only in the cortical gray matter, thalamus and striatum (Fig. 5B, Table 5). Similarly, we find significant correlations between local densities of endothelial cells and glial cells within some structures, but no strong systematic correlation across locations (Fig. 5C).

Consistently with the larger variation of neuronal densities than glial cell densities across locations, the glia/neuron (G/N) ratio is highly variable within the mouse brain, and is universally and inversely related to local neuronal densities across all sites and structures (Fig. 6A, B). That is, there are more glial cells per neuron in those sites with lower neuronal densities, in a relationship that applies both across and within structures, regardless of their identity (Table S5). In contrast, the relationship between local glia/neuron ratio and glial cell density is specific to each structure, with different ranges of glia/neuron ratios sharing similar glial cell densities across structures (Fig. 6C, D). Importantly, although higher G/N ratios where neuronal densities are smaller (and therefore neurons are larger) have been proposed to accompany the presumably higher energetic demands of larger neurons (Attwell et al., 2001), the entire 10,000-fold range of G/N ratios occurs within the same restricted range of endothelial cell densities, with no consistent correlation across brain structures and sites (Figure 6E, F). That is, endothelial cell densities are not significantly higher in those locations where each neuron is accompanied by a larger number of glial cells.

**Figure 6.**
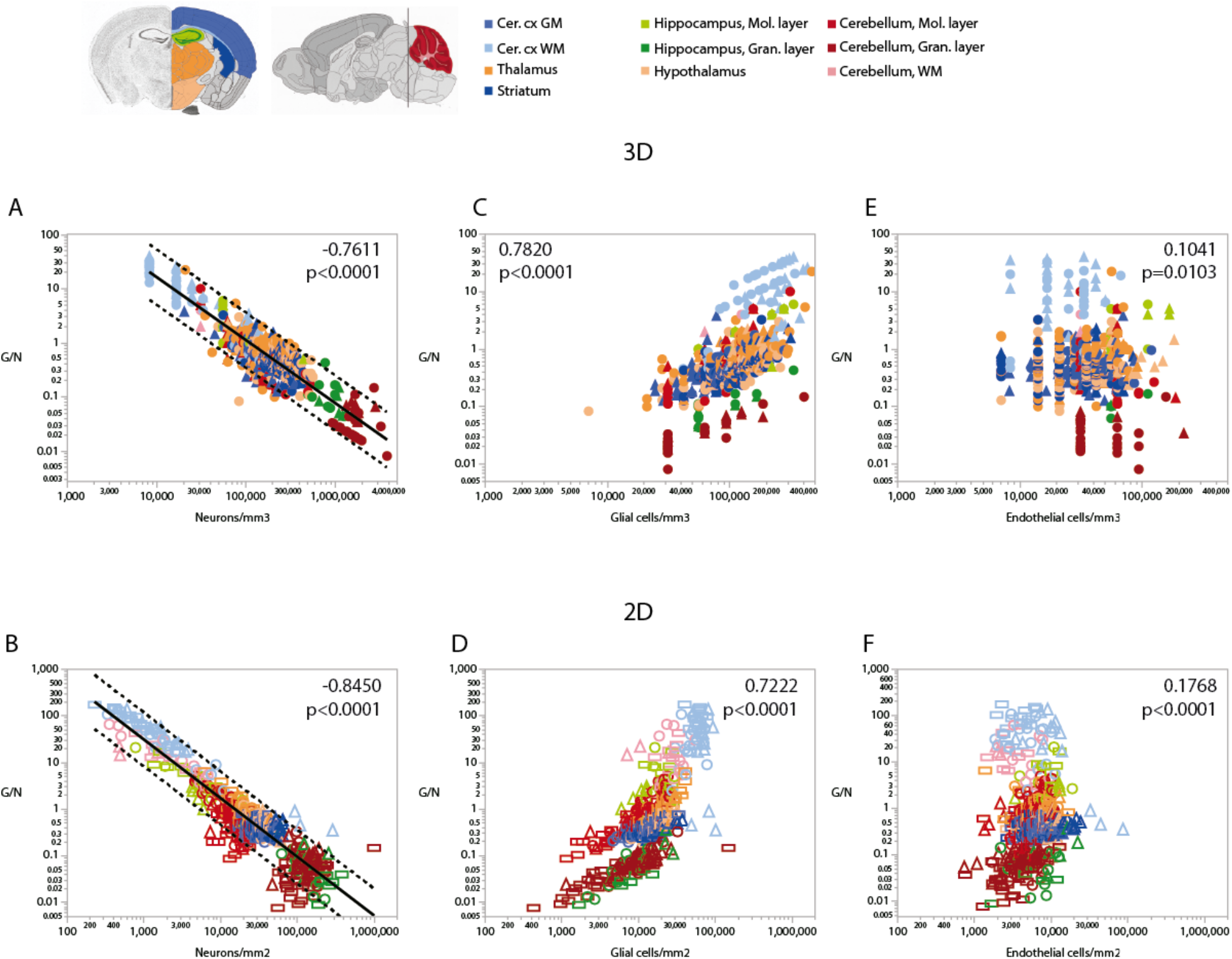
Ratio of glial cells per neuron (G/N) increases across brain structures and sites with decreasing neuronal densities. Spearman correlation coefficients and p-values across all structures and sites are indicated in each graph; relationships for each structure are given in Table 5. G/N ratios span over three orders of magnitude and vary as a single power function of neuronal density across all brain structures and sites whether measured in 3D stacks (**A**) or in 2D composite images **(B).** Power functions are plotted with 95% confidence interval (dashed lines), and have exponents −1.162±0.024, r^2^=0.816, p<0.0001 (**A**, 3D) or −1.257±0.020, r^2^=0.865, p<0.0001 (**B**, 2D). Notice that most data points in all structures are contained within the 95% CI of the function that applies across structures, indicating that the relationship is brain-wide. **C, D:** G/N ratio also correlates with glial cell densities across all brain structures and sites, but the relationships are clearly distinct across sites (see exponents in Table 5). **E, F:** Although a weak correlation is detected between G/N ratios and endothelial cell densities across brain structures and sites, most local correlations are non-significant or weak, with a wide range of non-overlapping values of G/N for similar endothelial cell densities across brain structures. All graphs are shown with similar X and Y scales for comparison.

Because of the much larger variation in neuronal densities than in endothelial cell densities across sites, the local ratio of endothelial cells per neuron is inversely proportional to local neuronal density across all sites and structures examined (Fig. 7). Importantly, this relationship is universal across brain locations, as the power function that applies across structures contains all structures and almost all data points within its 95% CI (Fig. 7A, B). That is, each neuron is accompanied by more endothelial cells at sites with lower neuronal densities than at sites with higher neuronal densities. In contrast, the ratio of endothelial cells per neuron does not vary consistently with local glial cell density neither across nor within brain structures (Fig. 7C, D). As expected for the quantity in the numerator of the ratio, local endothelial cell density is very strongly correlated with the E/N ratio in each brain structure in the mouse brain, but there is hardly any overlap across structures: because of the widely different neuronal densities across structures, brain sites with similar densities of endothelial cells have very different E/N ratios across structures (Fig. 7E, F). As could be expected of two quantities that are heavily dependent on variation in neuronal density, the E/N ratio is strongly and linearly correlated with the G/N ratio across all sites and structures (Fig. 8). That is, where more endothelial cells supply energy per neuron (i.e. in structure and sites with low neuron density), there are also more glial cells per neuron. Importantly, the relationship that applies across structures includes most data points for every structure, indicating that this relationship is universal.

**Figure 7.**
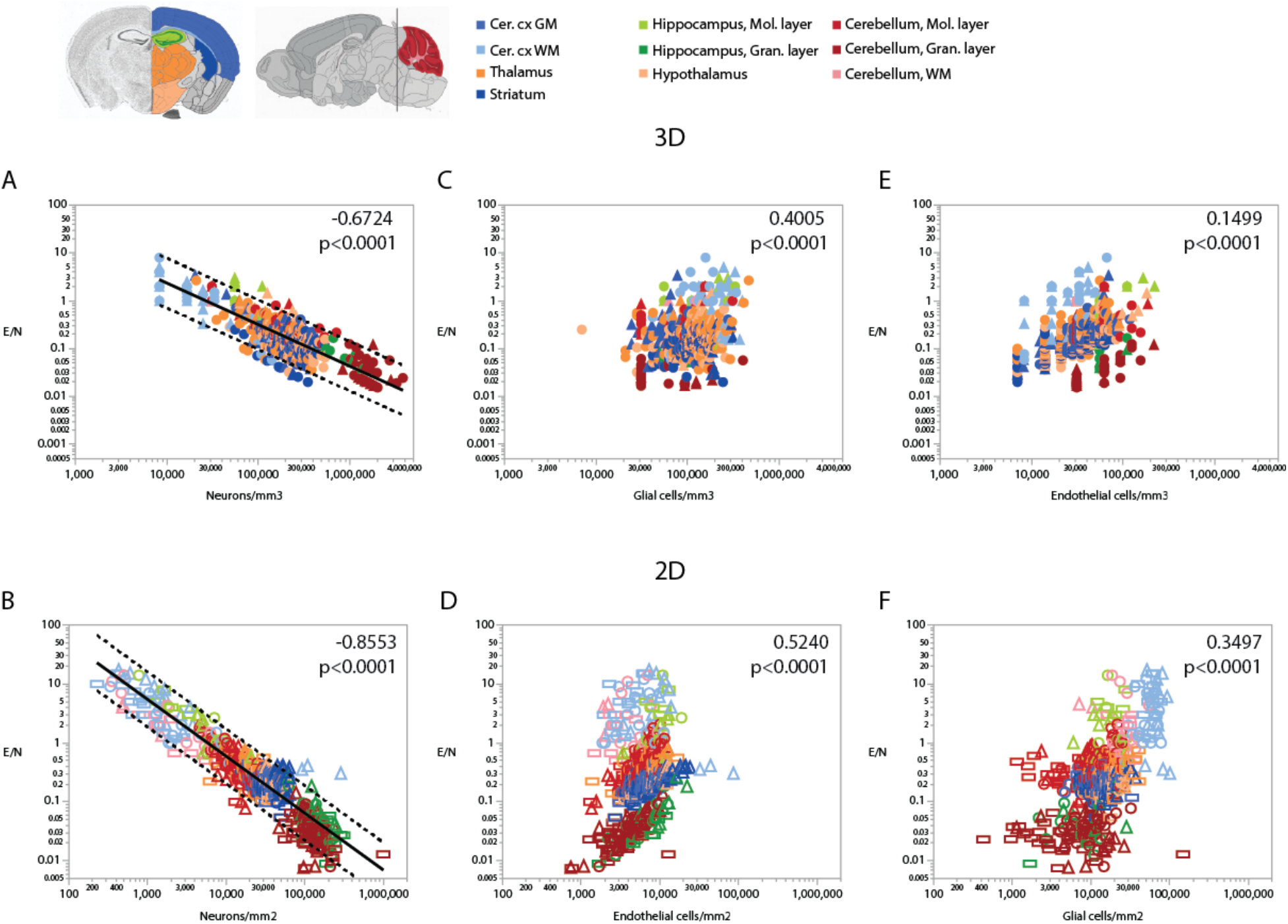
Ratio of endothelial cells per neuron (E/N) varies widely and uniformly across brain structures and sites depending on local neuronal density, but correlations with glial and endothelial cell densities only apply locally. Spearman correlation coefficients and p-values across all structures and sites are indicated in each graph; correlations and power function exponents for each structure are listed in Table 4. **A, B:** E/N ratios span three orders of magnitude across brain sites and structures and vary as a single power function of local neuronal density whether measured in 3D stacks (**A**) or in 2D composite images (**B**). **C, D:** E/N ratio appears to correlate with local glial cell density across brain structures in both datasets, but closer inspection reveals that the overall correlation results from the combination of locations and does not hold within most structures (see Table 5). **E, F:** In contrast, E/N ratios are strongly correlated with endothelial cell densities locally within each structure, but different brain structures have non-overlapping E/N ratios with similar endothelial cell densities, indicating that the apparent overall correlation does not result from a universal correlation, unlike that seen for the strong and universal correlation between E/N and neuronal density (**A, B**). All graphs are shown with similar X and Y scales for comparison.

**Figure 8.**
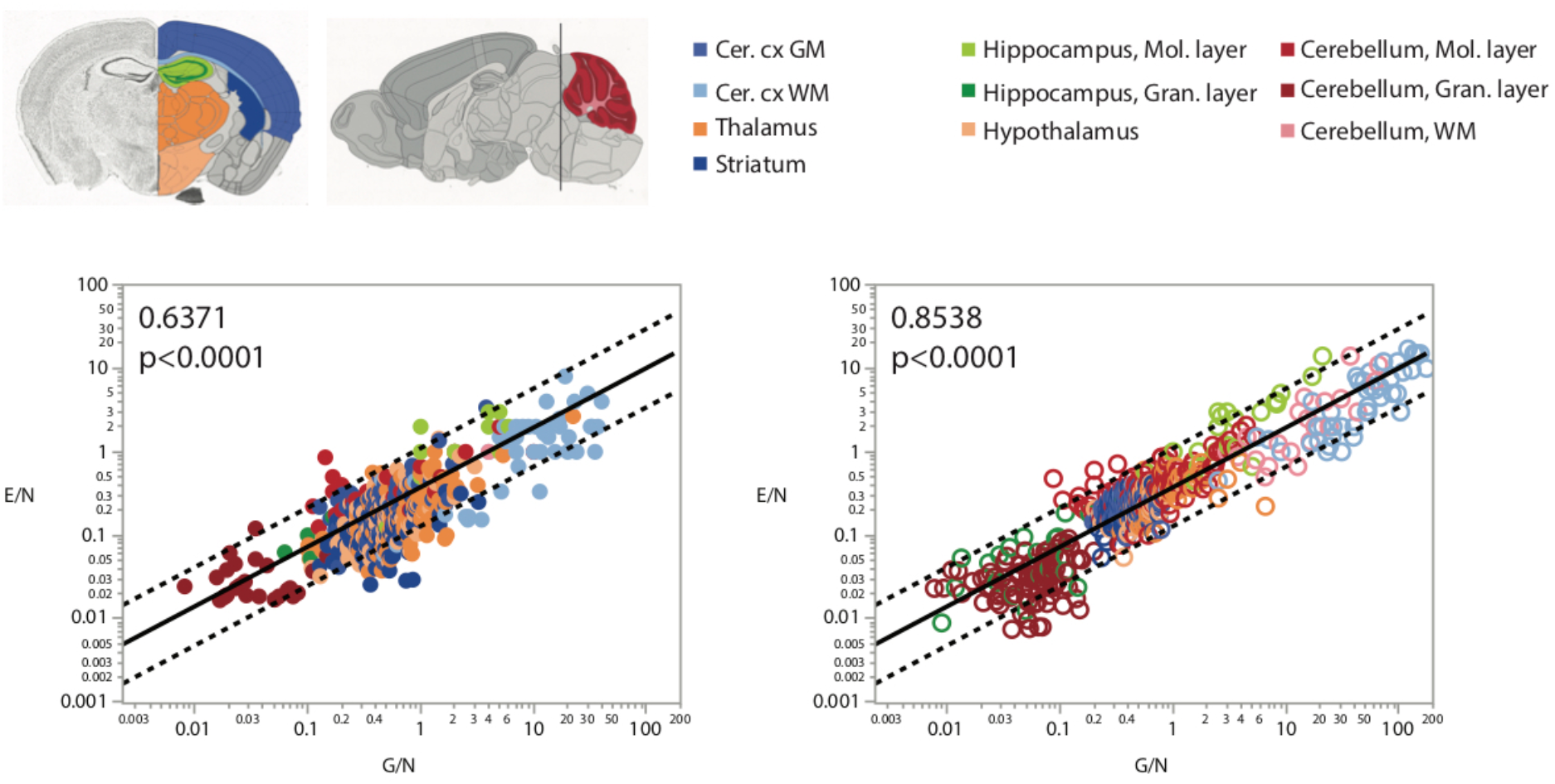
More endothelial cells per neuron (E/N) are uniformly found in brain structures and sites with more glial cells per neuron, as expected given that both these variables are strongly and inversely correlated with local neuronal density (Figs. 4, 5). Spearman correlation coefficients are indicated in the graphs.

Finally, comparing our measurements of local cell densities and capillary fraction in cortical grey matter sites matching those with recently published data on local synaptic densities in the mouse cerebral cortex (Zhu et al., 2018) we could examine whether there is evidence of higher energy availability in sites of higher synaptic densities to support the expectation that energy supply responds local demands due to synaptic activity (Attwell and Laughlin, 2001; Harris et al., 2012; Table 6). Densities of synapses (measured as synapses that express PSD and/or SAP according to data in Zhu et al., 2018) vary 1.5-fold across cortical sites, while densities of endothelial cells vary 2.5-fold and neuronal densities vary a similar 2.2-fold. However, we find no significant correlation between local endothelial cell density and synaptic density (Fig. 9A) or neuronal density (Fig. 9B) within the mouse cerebral cortex (Spearman, p=0.4133 and 0.7047), indicating that cortical sites with more synapses or more neurons receive as much capillary supply as sites with fewer synapses or neurons. Higher local synaptic densities are modestly associated with higher local neuronal densities (Fig. 9C; Spearman, p=0.0414), such that the ratio of synapses per neuron is concentrated in the 6,500-9,500 range (Fig. 7E; Spearman, p= 0.0129 due to a single data point above 12,500), compatible with previous estimates of ca. 8,000 synapses per cortical neuron in the mouse (Schüz, and Palm, 1989; Braitenberg and Schüz, 1998). We find that the average number of synapses per neuron is not higher where there are more synapses (Fig. 9D; Spearman, p=0.0775), and is also not significantly correlated with the local number of glial cells per neuron (Spearman, p=0.3177; Table 6). Indeed, local densities of synapses are not correlated with local glial cell densities (Spearman, p=0.6488), and the higher the local glial cell density, the fewer the synapses per glial cell (Spearman rho=−0.831, p<0.0001; Table 6). Crucially, the average number of synapses per neuron is not significantly correlated with the local ratio of endothelial cells per neuron (Fig. 9F; Spearman, p=0.4717): the E/N ratio varies about 4-fold across cortical sites with similar average numbers of synapses per neuron, and at a similar capillary supply per neuron, local neurons may have more or fewer synapses on average. Instead, the most striking pattern is, again, that at cortical sites where each neuron has more capillary cells available to it, there are also more capillary cells available per synapse (Fig. 9G; Spearman, p=0.0001).

**Table 6.** Spearman correlation coefficients and p-values for correlations across cerebral cortical sites. DE, local density of endothelial cells; DN, local neuronal density; DG, local density of glial cells; DS, local density of synapses; E/N, ratio of endothelial cells per neuron; E/G, ratio of endothelial cells per glial cell; E/S, ratio of endothelial cells per synapse; G/N, ratio of glial cells per neuron; S/N, ratio of synapses per neuron; S/G, ratio of synapses per glial cell. All values pertaining to DE, DN and DG were obtained in the present study, from analysis of 2D images acquired in locations matching those described in (*21*); all values pertaining do DS were calculated from (*21*).

|  | D<sub>E</sub> | D<sub>N</sub> | D<sub>G</sub> | D<sub>S</sub> | E/N | E/G | E/S | G/N | S/N | S/G |
| --- | --- | --- | --- | --- | --- | --- | --- | --- | --- | --- |
| D<sub>E</sub> |  | -0.084,<br>0.7047 | 0.412,<br>0.0509 | -0.194,<br>0.4133 | **0.753,**<br>**<0.0001** | **0.741,**<br>**<0.0001** | **0.906,**<br>**<0.0001** | 0.123,<br>0.5758 | -0.166,<br>0.4837 | -0.393,<br>0.0862 |
| D<sub>N</sub> | -0.084,<br>0.7047 |  | 0.307,<br>0.1542 | 0.460,<br>0.0414 | **-0.652,**<br>**0.0007** | -0.121,<br>0.5820 | -0.235,<br>0.3177 | **-0.624,**<br>**0.0015** | **-0.545,**<br>**0.0129** | 0.056,<br>0.8113 |
| D<sub>G</sub> | 0.412,<br>0.0509 | 0.307,<br>0.1542 |  | 0.108,<br>0.6488 | 0.128,<br>0.5605 | -0.203,<br>0.3525 | 0.312,<br>0.1803 | 0.441,<br>0.0350 | -0.291,<br>0.2131 | **-0.831,**<br>**<0.0001** |
| D<sub>S</sub> | -0.194,<br>0.4133 | 0.460,<br>0.0414 | 0.108,<br>0.6488 |  | -0.441,<br>0.0517 | -0.215,<br>0.3632 | **-0.506,**<br>**0.0230** | -0.306,<br>0.1896 | 0.404,<br>0.0775 | 0.363,<br>0.1156 |
| E/N | **0.753,**<br>**<0.0001** | **-0.652,**<br>**0.0007** | 0.128,<br>0.5605 | -0.441,<br>0.0517 |  | **0.630,**<br>**0.0013** | **0.750,**<br>**0.0001** | **0.516,**<br>**0.0118** | 0.171,<br>0.4717 | -0.306,<br>0.1893 |
| E/G | **0.741,**<br>**<0.0001** | -0.121,<br>0.5820 | -0.203,<br>0.3525 | -0.215,<br>0.3632 | **0.630,**<br>**0.0013** |  | **0.628,**<br>**0.0030** | -0.273,<br>0.2069 | 0.435,<br>0.0555 | 0.252,<br>0.2838 |
| E/S | **0.906,**<br>**<0.0001** | -0.235,<br>0.3177 | 0.312,<br>0.1803 | **-0.506,**<br>**0.0230** | **0.750,**<br>**0.0001** | **0.628,**<br>**0.0030** |  | 0.250,<br>0.2868 | -0.364,<br>0.1163 | **-0.526,**<br>**0.0171** |
| G/N | 0.123,<br>0.5758 | **-0.624,**<br>**0.0015** | 0.441,<br>0.0350 | -0.306,<br>0.1896 | **0.516,**<br>**0.0118** | -0.273,<br>0.2069 | 0.250,<br>0.2868 |  | 0.235,<br>0.3177 | **-0.669,**<br>**0.0013** |
| S/N | -0.166,<br>0.4837 | **-0.545,**<br>**0.0129** | -0.291,<br>0.2131 | 0.404,<br>0.0775 | 0.171,<br>0.4717 | 0.435,<br>0.0555 | -0.364,<br>0.1163 | 0.235,<br>0.3177 |  | 0.435,<br>0.0555 |
| S/G | -0.393,<br>0.0862 | 0.056,<br>0.8113 | **-0.831,**<br>**<0.0001** | 0.363,<br>0.1156 | -0.306,<br>0.1893 | 0.252,<br>0.2838 | **-0.526,**<br>**0.0171** | **-0.669,**<br>**0.0013** | 0.435,<br>0.0555 |  |

**Figure 9.**
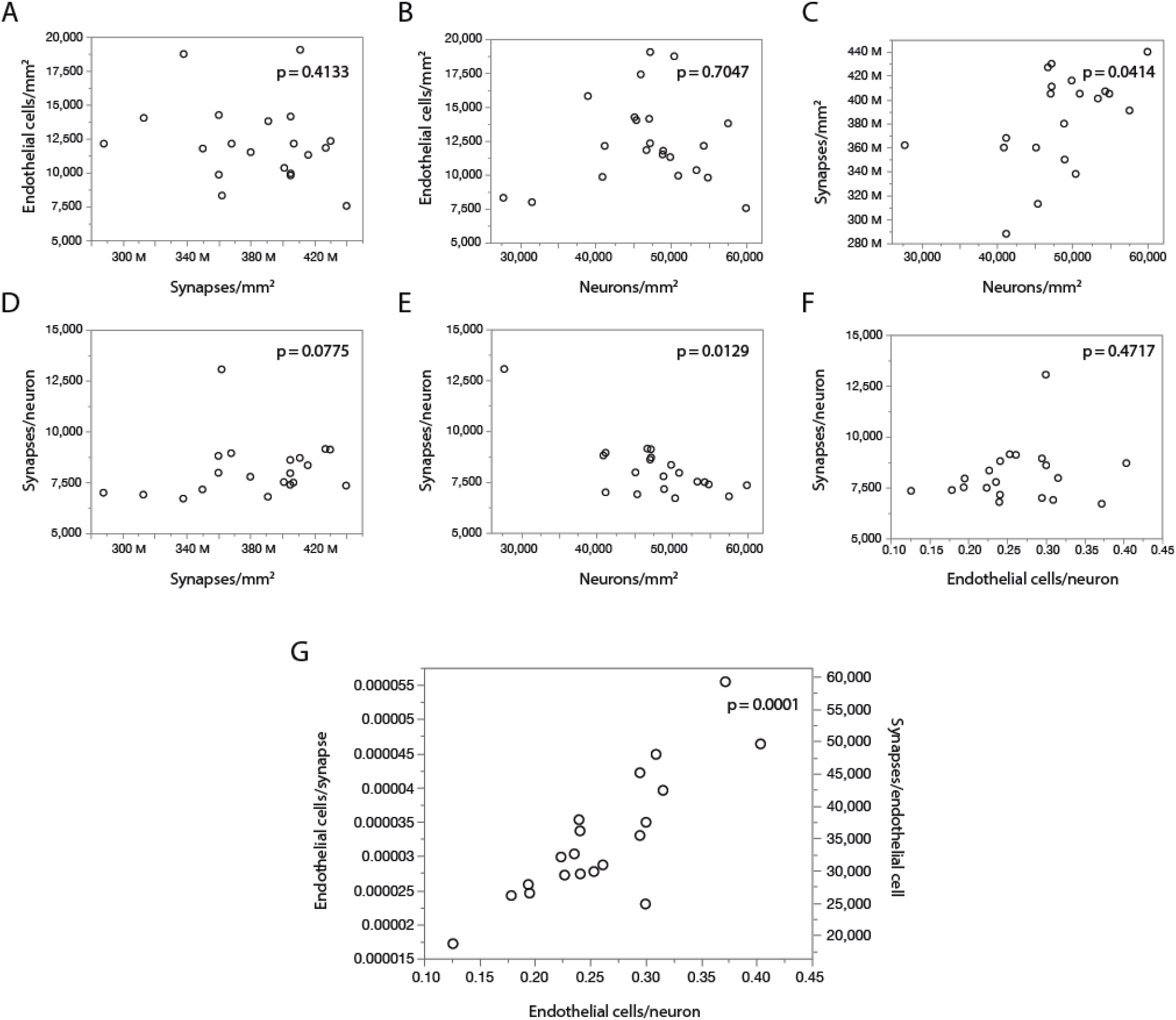
With fairly constant numbers of synapses per neuron, the number of synapses served per endothelial cell increases together with the ratio of endothelial cells per neuron across sites in the mouse cerebral cortex. Data on local synaptic densities from (Zhu et al., 2018) were combined to estimates of local densities of endothelial cells, neurons and glial cells in 2D images acquired in matching locations of the cerebral cortex as described in (Zhu et al., 2018). Local densities of endothelial cells are not significantly correlated with local synaptic densities (**A**) or neuronal densities (**B**). Sites with more neurons have marginally, but significantly, more synapses (**C**), which is consistent with a lack of correlation between local synaptic density and the number of synapses per neuron (**D**), although a correlation between average number of synapses per neuron and local neuronal density, due to one outlier, cannot be discarded (**E**). Still, there is very clearly no correlation between local numbers of synapses per neuron and the ratio of endothelial cells per neuron (**F**); rather, the latter is directly, and strongly (r^2^=0.750), correlated with the ratio of endothelial cells per synapse (or its inverse, the number of synapses per endothelial cell; **G**). All correlation coefficients are available in Table 6.

These data thus indicate that higher E/S is associated with higher E/N, even though there are not more endothelial cells where there are more synapses (or more neurons; Fig. 9A, B), and there are also not more E/N where there are more S/N (Fig. 9F). Rather, higher E/S occurs with higher E/N (Fig. 9G) simply due to small variations in endothelial cell densities that are not correlated with variations in densities of neurons or synapses (Fig. 9A-B) in the presence of significant local variations in neuronal densities that are only somewhat correlated with smaller variations in densities of synapses (Fig. 7C). Principal component analysis supports this conclusion: three factors are required to account for the variance in all six variables, with capillary density loading only in the first factor (explaining 42.9% of variance; factor loading, 0.996, associated with E/N [0.816] and E/S [0.926]), neuronal density loading only in the second factor (total variance explained, 78.8%; loading, −0.943, associated with E/N [0.557] and S/N [0.976]), and synaptic density as the sole contributor to the third factor (explaining a total of 99.3% of variance; factor loading, 0.981).

Finally, while the glia/neuron ratio is locally higher in cortical sites where the endothelial cell/neuron ratio is also higher (Spearman, p=0.0118), the number of endothelial cells per synapse is not correlated with the local glia/neuron ratio (p=0.2868; Table 6). Thus, a higher proportion of glial cells per neuron is not associated with a larger capillary supply per synapse.

## Discussion

While neuroscientists often frame the energy *cost* of the brain in terms of high neuronal *demand* for energy, we prefer to reframe metabolism simply as energy *use*, a term that implies nothing about what defines it, whether supply or demand, and instead leaves these open to investigation. The issue of energetic *use* by neurons can then be separated into the more tractable questions of (a) whether there is evidence that larger neurons *demand* and thus are provided with more energy, depending on their intrinsic biophysical properties (Attwell and Laughlin, 2001), and (b) whether there is evidence that larger neurons are *supplied* with more energy than smaller neurons, regardless of (a). Our finding that local capillary area fraction or density of capillary cells does not accompany the enormous variation in neuronal densities across sites in the mouse brain nor synaptic densities in the mouse cerebral cortex provides evidence against (a) and in support of (b), against several intuitive, but so far untested, central tenets of neurophysiology and functional brain imaging, namely: that neurons *demand* energy, and larger neurons *demand* more energy than smaller neurons; that sites with more synapses use more energy; that because larger neurons have more synapses, they demand more energy; and that the steady-state density of the adult capillary bed reflects self-organized adjustments across brain sites according to variations in local energy demand. Instead, we interpret our data to indicate that, due to the 100x larger variation in neuronal densities than in capillary density throughout the mouse brain, larger neurons *get access* to more energy simply because where neurons are larger, as indicated by lower neuronal densities, fewer neurons compete for the capillary density-restricted blood supply. Similarly, we propose that, given a small but significant variation in densities of synapses with densities of neurons within the cerebral cortex, there is a restricted range of numbers of synapses per neuron, and more endothelial cells providing energy both per neuron and per synapse where neurons are larger (highlighted middle panel in Fig. 1). These findings suggest the intriguing possibility that increasing neuronal sizes in evolution may have brought neurons the advantage of not decreasing energy availability per cell, and possibly holding that constant, as body size increases and total metabolic rate scales more slowly than body mass (Herculano-Houzel, 2011), a possibility that we are now ready to address by extending the present analysis to other species.

Within the cerebral cortex in particular, a previous detailed study of variation in fractional volume of the microvasculature across layers found that it varies little, and does not reflect variations in neuronal density across cortical layers (Tsai et al., 2009). When compiling data across layers, that study did find a significant positive correlation between neuronal density and microvasculature volume fraction across sites, which our analysis corroborates when restricted to the cerebral cortical gray matter alone (Table 5). Importantly, while this variation of capillary density with neuronal density is significant across cortical sites, it is by no means large enough to maintain a constant E/N ratio across cerebral cortical sites within the mouse brain: even within the cerebral cortical gray matter, the E/N ratio still decreases with increasing neuronal density (Table 5), which is indicative of less energy availability per neuron in these sites. Therefore, it remains true for the cerebral cortex alone that our data suggest that energy availability per neuron is supply-limited.

There are presently no direct data available on rates of energy use per neuron, whether in different structures or species, to test the hypothesis that larger neurons use more energy (whether due to supply or demand); all measurements so far have been of energy use per gram of tissue (Mink et al., 1981; Karbowski, 2007), not per neuron. A necessary first step to establishing whether larger neurons *use* more energy, be it due to increased demand, supply, or both, is comparing average energy use per neuron where neurons have different sizes, both within and across species. The practical impediment here is that measuring energy use requires bringing live animals to the lab. We propose that local capillary density, which correlates very well, and linearly, with resting blood flow and local rate of glucose use in both rat (Klein et al., 1986; Borowsky and Collins, 1989) and macaque brains (Noda et al., 2002), and can be measured readily and efficiently in 2D images of thin sections of fixed brains, is a good proxy and viable alternative to estimating energy availability per volume as well as per neuron in fixed brain tissue. While direct validation of the correspondence between local capillary density and resting metabolic rate is underway, we expect that systematic analyses of local capillary densities will provide a powerful tool to circumvent the limitations to studying energy use per neuron in living animals. Indeed, we find that the small variations in the local vascular fraction measured as in the present study across sites in the rat brain are a very good approximation of measurements of local energy cost at rest in each structure (Ventura-Antunes, Dasgupta and Herculano-Houzel, in preparation). These findings open the way to comparative studies of energy availability per neuron in brains of species that are unlikely to ever be brought alive to laboratory settings.

While we examine fixed tissue and can make no claims about physiological properties and dynamic changes in neurovascular coupling (Leithner and Royl, 2014), the fairly stable distribution of the capillary bed in the perfused tissue compared to neuronal densities does inform that there is more energy available per neuron in those sites with lower neuronal densities (with few and larger neurons; Mota and Herculano-Houzel, 2014) than in sites of high neuronal density. These findings are compatible with a scenario where neurons compete for a limited energy supply, rather than the usually presumed scenario in which larger neurons and/or neurons with more synapses *demand* more energy which is then provided as the capillary bed adapts to those demands during development. A limited energy supply constrained by the density of the capillary bed would have direct consequences for the level of neuronal and synaptic activity that can be sustained across brain sites, in line with our previous suggestion that fundamental aspects of neuronal structure and function are constrained by energy supply across species (Herculano-Houzel, 2011). It remains to be determined whether these constraints manifest themselves during development, for instance through the self-regulation of the numbers of synapses that can form and remain active, or simply through self-regulation of levels of activity (as in synaptic homeostasis; Turrigiano, 2012).

The present findings have fundamental implications for brain health and normal and diseased aging. First, they suggest that the heightened vulnerability of the human white matter to ischemia (Wang et al., 2016) may be primarily due to its low capillary density compared to gray matter structures, and not simply to its large relative volume. Second, as cortical locations with high neuron densities have far less blood supplied per neuron than locations with low neuronal densities, individual neurons in “crowded” cortical areas are likely to be much more vulnerable to aging and pathologies that compromise circulation and/or metabolic capacity. Most remarkably, the hippocampus, whose granular layer contains most of its neurons and exhibits some of the lowest values of E/N found here, is one of the brain structures most highly vulnerable to hypoxia (Hossmann, 1999). Finally, some of the regions first and most vulnerable to Alzheimer’s disease are cortical areas with high resting metabolism (Buckner et al., 2005). While that might be indicative of higher capillary densities that would increase energy availability as a whole, it is possible that this is offset by very high neuronal densities, such as those found in the entorhinal cortex and hippocampus here, leading to low values of E/N. We are currently investigating the possibility that local variation in E/N correlates with brain tissue vulnerability in normal and diseased aging.

## Acknowledgments

Thanks to Louise Botelho, Felipe Tenório and Marina Ricardo for contributing to image analysis, and to Prof. Doug Rothman for insightful discussions. This work was supported by a Scholar Award from the James S. McDonnell Foundation and Vanderbilt start-up funds to SHH, and by a CAPES fellowship to LVA. SHH conceived the study, LVA executed all data acquisition, and both authors analyzed data and wrote the manuscript.

